# Dissecting muscle synergies in the task space

**DOI:** 10.1101/2023.03.17.533096

**Authors:** David Ó’ Reilly, Ioannis Delis

## Abstract

The muscle synergy is a guiding concept in motor control research that relies on the general notion of muscles ‘*working together’* towards task performance. However, although the synergy concept has provided valuable insights into motor coordination, muscle interactions have not been fully characterised with respect to task performance. Here, we address this research gap by proposing a novel perspective to the muscle synergy that assigns specific functional roles to muscle couplings by characterising their task-relevance. Our novel perspective provides nuance to the muscle synergy concept, demonstrating how muscular interactions can ‘*work together’* in different ways: a) irrespective of the task at hand but also b) redundantly or c) complementarily towards common task-goals. To establish this perspective, we leverage information- and network-theory and dimensionality reduction methods to include discrete and continuous task parameters directly during muscle synergy extraction. Specifically, we introduce co-information as a measure of the task relevance of muscle interactions and use it to categorise such interactions as task-irrelevant (present across tasks), redundant (shared task information) or synergistic (different task information). To demonstrate these types of interactions in real data, we firstly apply the framework in a simple way, revealing its added functional and physiological relevance with respect to current approaches. We then apply the framework to large-scale datasets and extract generalizable and scale-invariant representations consisting of subnetworks of synchronised muscle couplings and distinct temporal patterns. The representations effectively capture the functional interplay between task end-goals and biomechanical affordances and the concurrent processing of functionally similar and complementary task information. The proposed framework unifies the capabilities of current approaches in capturing distinct motor features while providing novel insights and research opportunities through a nuanced perspective to the muscle synergy.

## Introduction

Human movement is a highly complex behaviour, with a broad spectrum of multiplexed spatiotemporal dynamics typically exhibited for basic activities-of-daily-living [1,2]. How the central nervous system controls movement in the face of this inherent complexity to ensure efficient and reliable navigation of the environment and task performance is a nontrivial question currently under investigation in the motor control field [3,4]. The muscle synergy hypothesis is a long-withstanding proposition on the underlying neural constraints producing coordinated movement, stating that this complexity is offset by the allocation of computational resources to the spinal-level in the form of motor primitives [3–6]. These motor primitives modularly activate functional groups of muscles that are flexibly combined for the efficient construction of a movement. The conceptual underpinning of the ‘*muscle synergy’* entails the idea of combinations of muscles ‘*working together’* for the purpose of effective goal-orientated behaviour [7]. This emergent cohesion involves the following qualifying attributes: a repeatable muscle activation pattern common across trials and participants, a reciprocal relationship among functional muscle groups such that changes occur in one group to compensate for changes in another, and task dependence (i.e. the pattern of interdependencies among muscles must map onto task performance) [7,8]. A common approach to analyse the neural constraints underlying these motor patterns is to apply unsupervised machine-learning algorithms to electromyographic (EMG) data [9,10], with the aim of extracting a latent, low-dimensional representation.

In [11], we considered, key limitations among current approaches to muscle synergy analysis in extracting functionally relevant and interpretable patterns of muscle activity [12]. We proposed a combinatorial approach based on information- and network-theory and dimensionality reduction (the network-information framework (NIF)) that significantly improved the generalisability of the extraction process by, among others, removing restrictive model assumptions (e.g. linearity, same mixing coefficients) and the reliance on variance-accounted-for (VAF) metrics [12]. By determining the pairwise mutual information between muscles, this innovation paved the way for the appropriate mapping of muscular interactions to the task space. To elaborate on the significance of this development, the extraction of motor patterns in isolation of the task space often comes at the expense of functional and physiological relevance [12,13]. Furthermore, effective methods for mapping large-scale physiological dynamics to behaviour is a current gap across the neurosciences [14]. Thus, here we build on this work by, for the first time, directly including task space parameters during muscle synergy extraction. This enables us, in a novel way, to dissect the concept of the muscle synergy and therefore quantify interactions between muscle activations with shared or complementary functional roles.

Further to the above, in its currently defined state, the muscle synergy concept describes the role of common neural drives to functional muscle groupings working redundantly towards a common task-goal [3–5,7,15]. However, recent influential works have highlighted several other important mechanisms involved in this low-dimensional control strategy that are not well recognised by the muscle synergy concept [16–24]. Such insights include the partitioning of motor variability by the nervous system into task-relevant and -irrelevant spaces and the cooperation between functionally distinct muscle groupings in the form of cross-module functional connectivities. These observations highlight the need for a refinement of the muscle synergy concept to comprehensively describe diverse muscle interactions during movement, including their partitioning into task-relevant and –irrelevant spaces and the characterisation of their functional roles.

We thus motivate the development of a more nuanced perspective to the muscle synergy concept and the general notion of ‘*working together’* that comprehensively describes the muscle interactions underlying motor behaviour. To do so, we propose an information-theoretic approach (based on the NIF pipeline) that characterises the contributions of muscle couplings to task performance. In other words, we frame the notion of ‘*working together’* in more specific terms of shared information between pairs of spatiotemporal muscle activations ([*m*_*x*_, *m*_*y*_] (Fig.1.3(a))) (red and green sets respectively) and a corresponding task parameter (*τ*) (blue set) (see Venn diagrams in Fig.1.3(b) below). Among current approaches to muscle synergy analysis, the shared information between muscles (yellow and white area in Fig.1.1(a)) is quantified, using dimensionality reduction, as common patterns of variability. These common patterns are essentially task-agnostic and may contain patterns of variability a) present in specific tasks (i.e. task-relevant (white shaded area in Fig.1.1(a)) as well as b) shared across tasks (i.e. task–irrelevant (yellow shaded area in Fig.1.1(a)). Our proposed approach dissects patterns of muscle variability in space, time and across trials in terms of their task-relevance and functional similarity in a generalizable manner using mutual information (MI) (Fig.1.3(a-c)). This enables us to decompose muscle activations into muscle pair – task parameter couplings and characterise their combined functional roles. We can then extract low-dimensional representations of these muscle couplings, i.e. muscle networks with specific spatial and temporal signatures, across participants and tasks (Fig.1.3(c)).

**Fig. 1:**
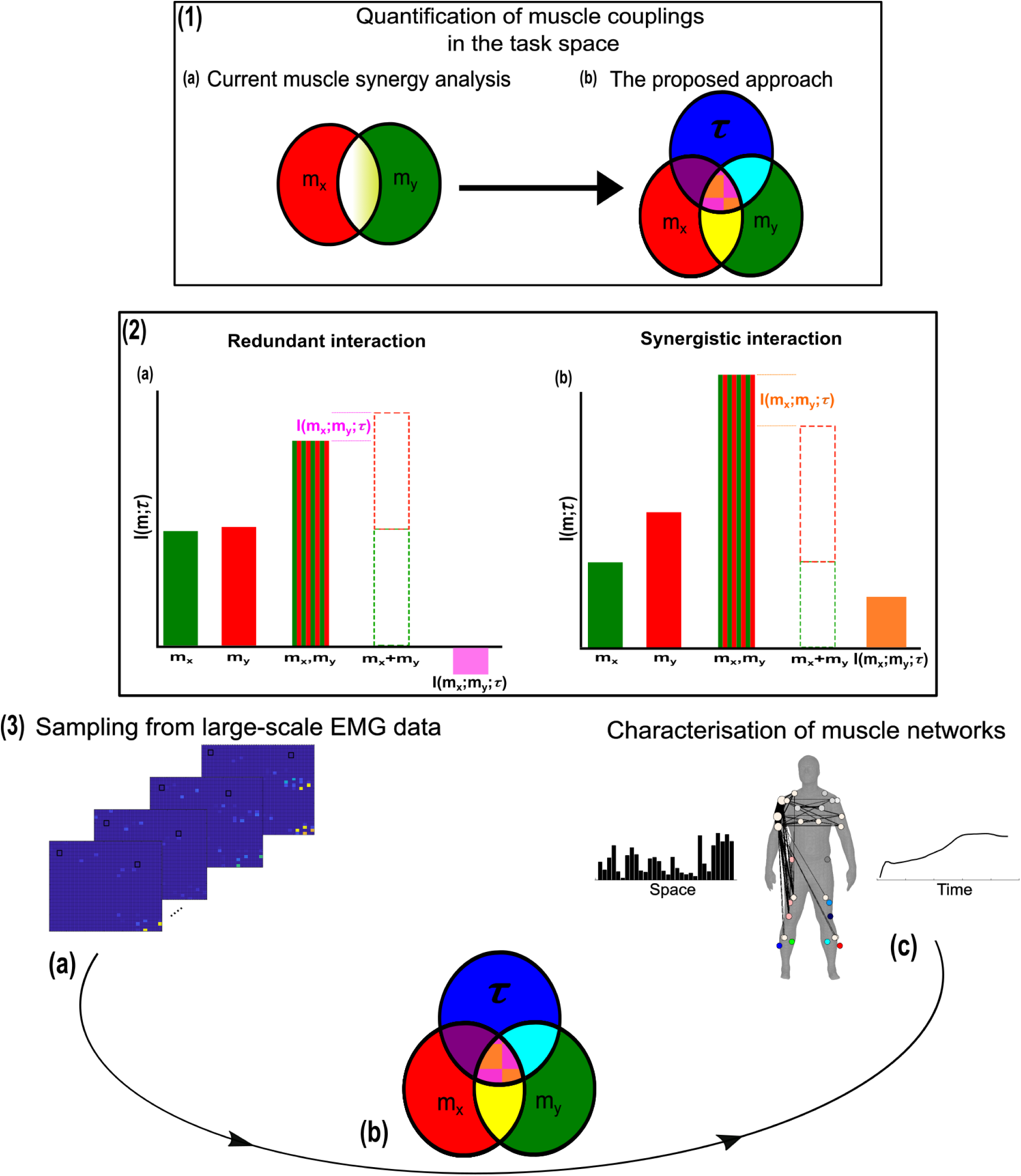
A general outline of the proposed approach. **(1.1(a-b))** We propose a novel approach to mapping muscle couplings to the task space. Among current muscle synergy analysis approaches, muscle couplings are quantified in isolation of the task solely using dimensionality reduction. Using our approach, the functional characteristics of muscle interactions can be quantified in terms of the similarity of their encoded task information. We do so by determining the coupling between [*m*_*x*_, *m*_*y*_] and a corresponding task parameter (*τ*) using mutual information (MI). From this perspective, task-redundant muscle couplings (pink shaded area in pink-orange intersection) represent muscles cooperating towards similar task goals, while task-synergistic muscle couplings (orange shaded area in pink-orange intersection) encapsulate the task information provided by a muscle pairing acting towards complementary task goals. Muscle couplings present across tasks (i.e., task-irrelevant) are quantified by conditioning the MI between [*m*_*x*_, *m*_*y*_] pairs with respect to *τ* (yellow intersection). **(1.2)** A description of redundant and synergistic interactions. **(a)** Net redundant interactions are defined by a greater amount of information generated by the sum of individual observation of *m*_*x*_ and *m*_*y*_ ([*m*_*x*_ + *m*_*y*_]) than their simultaneous observation ([*m*_*x*_, *m*_*y*_]). **(b)** In a net synergistic interaction, [*m*_*x*_, *m*_*y*_] provides more information than [*m*_*x*_ + *m*_*y*_]. **(1.3(a-c))** An overview of the approach. Spatiotemporal muscle activation samples are extracted across trials from large-scale EMG datasets and concatenated into vectors, forming [*m*_*x*_, *m*_*y*_] pairs.The derived muscle couplings are then run through the NIF pipeline [11], producing low-dimensional, multiplexed space-time muscle networks.

Crucially, using this novel framework, we can separately quantify the task-irrelevant (i.e. muscle interactions present across tasks) information conveyed by a muscle coupling (yellow intersection in Fig. 1.1(b)) from the task-relevant information (pink-and-orange shaded area in Fig.1.1(b)). These task-relevant interactions can be either sub-additive/redundant (i.e. the muscle coupling provides less information about the task compared to the sum of the individual muscle patterns) or super-additive/synergistic (i.e. the muscle coupling conveys more task information than the sum of individual muscle-task encodings). Conceptually, the information a muscle interaction provides is considered redundant when all the information can essentially be found in one of the muscles (pink shaded area (Fig.1.2(a)). This redundant task information thus reveals a functional similarity between muscle activations. Alternatively, we can also identify muscles that act synergistically towards complementary task goals, meaning their variations provide different information about a motor behaviour. A key, quantifiable attribute of this complementary interaction is the emergent task information (‘*synergy*’) they provide when considered together (orange shaded area (fig.1.2(b)). From this novel perspective, muscle activations can ‘*work together’* not just similarly towards a common task-goal but also complementarily towards different aspects of motor behaviour and concurrently towards objectives functionally irrelevant to overt task performance, thus providing a comprehensive view of the muscle interactions governing coordinated movement.

To illustrate this novel conceptual and analytical framework, we conducted two example applications to data from human participants performing naturalistic movements. These applications demonstrate the added utility of this framework to current muscle synergy analysis in terms of functional and physiological relevance and interpretability. We then applied it to three large-scale datasets, extracting generalizable and functionally interpretable space-time muscle networks with respect to both discrete and continuous task spaces. We have also made available open-source Matlab routines for readers to apply this approach to their own data (https://github.com/DelisLab/EMG2Task).

## Results

Our primary aim here is to characterise muscle synergies in task space by quantifying the contributions of muscle couplings to task performance. To achieve this, we essentially reverse the analytical approach typically used in muscle synergy studies (i.e. muscle groupings are identified and inferences then made about their functional roles) [9]. More specifically, we firstly identify functional couplings between paired muscle activations by evaluating their task-relevance and then extracting representative patterns of such couplings using dimensionality reduction methods. This enables us to distinguish task-irrelevant from task-relevant muscle couplings. Of the muscle couplings that demonstrate task-relevance, we can then characterise their functional roles as either redundant or synergistic. Fig.2 illustrates a simulation to facilitate interpretations of what informational redundancy and synergy mean when applied to muscle activities in the context of task performance. Redundant task information is generated when *m*_*x*_ and *m*_*y*_ carry identical predictive information about *τ*. This is distinct from current muscle synergy analysis which would consider *m*_*x*_ and *m*_*y*_ to share information about *τ* if their magnitudes are equivalent. Here, *τ* is always L when *m*_*x*_ is on and when *m*_*y*_ is off and R when vice versa. Thus, all the task information can be found in either *m*_*x*_ or *m*_*y*_ alone, generating 1 bit of redundant information. Synergistic task information on the other hand is predictive task information generated only when observing both *m*_*x*_ and *m*_*y*_ together. In the simple example shown, *τ* can be L when both *m*_*x*_ and *m*_*y*_ are active or inactive. However, we can see that when both muscles are active or inactive then *τ* is L. Thus, no predictive task information is provided by either *m*_*x*_ or *m*_*y*_ alone but the full 1 bit of information available is generated when observing both muscles together.

**Fig. 2:**
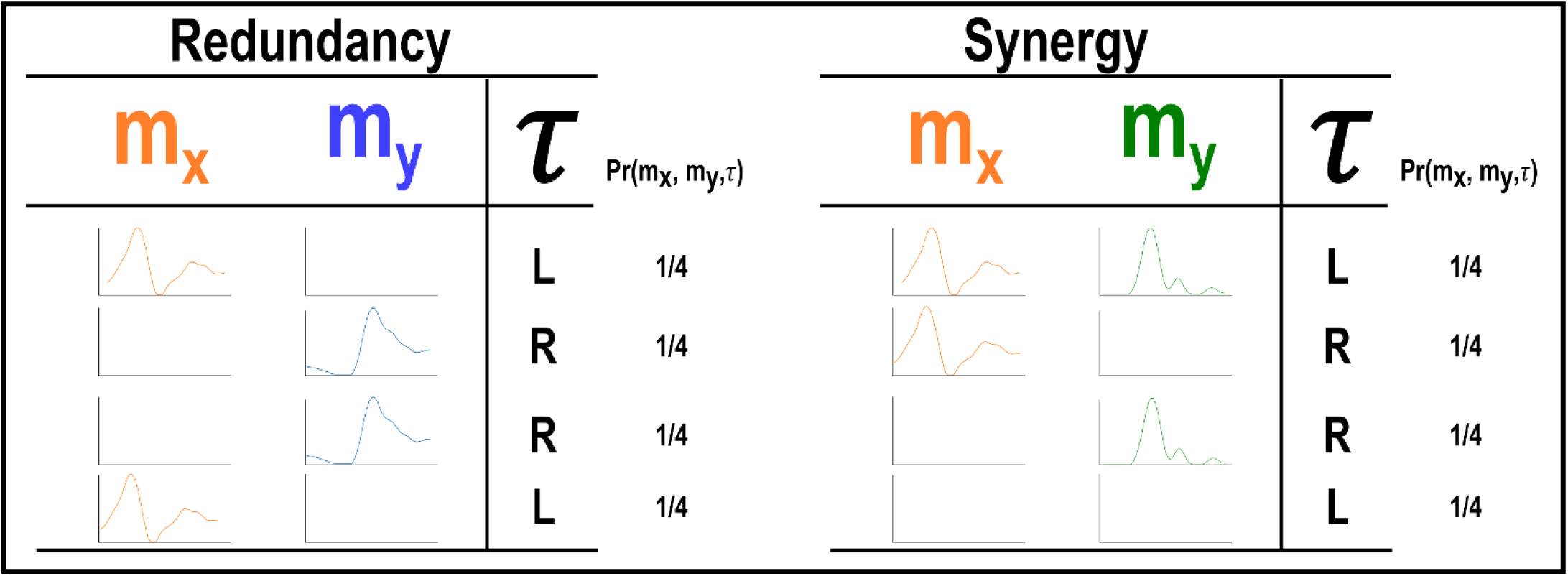
A simulation demonstrating how informational redundancy and synergy can be interpreted when applied to the muscle space. Four observations of a given muscle pair (*m*_*x*_ and *m*_*y*_) that can fall into two equiprobable on and off activation states and a corresponding task parameter (*τ*) describing left (L) or right (R) movement direction. Observing either *m*_*x*_ or *m*_*y*_ in the redundancy example gives 1 bit of information while observing both *m*_*x*_ and *m*_*y*_ together in the synergy example gives 1 bit of information.

### Building on current approaches to muscle synergy analysis

Current approaches to muscle synergy analysis based on non-negative matrix factorisation (NMF) have a proven use case in the extraction of functionally- and physiologically relevant motor patterns [3,12,34–36]. To demonstrate that the proposed framework adds to this current utility, here we firstly provide a simple example output from the proposed and current approaches (see fig.5(A-D)). This example was derived from the EMG recordings of a single-trial of a participant walking on level-ground in the counter-clockwise direction around the circuit depicted in fig.4(C) [27]. For the proposed approach, the muscle couplings were determined with respect to a single, continuous task parameter, the heel kinematic marker in the anterior-posterior direction. For the application of the current approach, we applied the spatial muscle synergy model across the same single-trial EMG recordings [37], extracting one component.

The simplified representations from the proposed approach reveal the functional role of muscular interactions with respect to the heel marker and provide intuition on the types of muscle couplings that can be identified, including task-irrelevant (A) -redundant (B) and –synergistic (C) interactions. Their submodular structure are illustrated via the node colour on the accompanying human body models [38], describing muscles that have a closer functional relationship. For example, a) the hamstring muscles (ST and BF) controlling knee flexion work redundantly together with muscles involved in hip abduction (GR, GlutM and Obl) to move the heel around the circuit (Fig.4(B)), and b) calf muscles involved in ankle flexion (GM, TA) cooperatively determine heel position in synergy with the same hamstring muscles (Fig.5(C)). Moreover, these hamstring muscles together with their antagonist RF also form a task-irrelevant network, i.e. their couplings are not predictive of heel position (Fig.5(A)). Overall, task-irrelevant muscle couplings primarily capture interactions between co-agonist and agonist-antagonist muscles that are indiscriminative of heel marker position. Also, task-redundant and -synergistic couplings reveal functionally similar and dissimilar muscle combinations respectively that provide sub-additive (i.e. shared or redundant) and super-additive (i.e. complementary or synergistic) information about heel marker position.

The NMF representation (fig.5(D)) conveys important information about gait in a task-agnostic manner, thus it may contain both task-relevant and -irrelevant interactions. Intuitively, SO, GM and Obl of the outer, right-side are most prominently weighted here, perhaps representing their functional role in accelerating the body in the counterclockwise direction around the circuit whilst maintaining upright posture (fig.4(C)). This observation can only be inferred indirectly however as, amongst current approaches, no direct association is made with other muscles or with task performance. Indeed, with respect to heel position, the proposed approach reveals that SO is functionally similar to Obl and GM whilst also containing task-synergistic and -irrelevant information with GM (fig.5(A-C)). From the perspective of existing approaches, the knee flexors and extensors play a minor role during turning gait (as indicated by their relatively low weighting (fig.5(D))). However, the proposed approach conveys a central role for these muscles suggesting that the proposed framework enables targeted dissections of muscle functionality with respect to a chosen task parameter, thus revealing subtle couplings with potentially important behavioural consequences.

Next, to demonstrate the additional physiological relevance the proposed methodology brings to current muscle synergy analysis, we applied the proposed framework to single trials of pointing movements performed by 20 participants with stroke and 25 healthy controls from dataset 4 (fig.4(D)) (‘MMH’ task 9 [33]). Specifically, we determined MI from a randomly selected trial of healthy and post-stroke participants with respect to the 3D position of the anterior wrist kinematic marker (WRBA) of the pointing arm. We chose WRBA as the task variable here due to its sensitivity to hand orientation. For the purpose of this simplified demonstration, we focused on a comparison between task-redundant (with respect to WRBA) and NMF-based muscle representations. We firstly generated a normative representation of this pointing motion by extracting the first component across the healthy controls using the proposed approach (Fig.6(a.1)) and NMF (Fig.6(a.2)). We then quantified the similarity of muscle representations extracted from each post-stroke participant individually to this normative reference (obtained across all healthy participants) using Pearson’s correlation and converted these values to distances (i.e. 1-r) (Fig.6(b-c)). Finally, we determined if these distances from healthy control values were predictive of the stroke survivor’s motor impairment, measured using the upper-extremity section of the Fugl-Meyer assessment (FMA-UE) (Fig.6(d)).

To briefly summarise the results, the distance of post-stroke participants from healthy controls was found to be predictive of motor impairment for the proposed approach (*β*= -8.52±2.2, p=0.0012) but not the NMF-based approach (*β*= 1.4±1.39, p=0.33). This finding suggests, intuitively, that the proposed approach captures redundant muscle couplings that support robust motor control and that deviations from this normative pattern of motor redundancy are linearly related to the degree of impairment. Importantly, this result was obtained using only one randomly selected trial for each participant. This simple example conveys the physiologically-relevant targeted insights that can be generated from the proposed framework. Although current approaches have demonstrated significant linear relationships with motor impairment [40–42], these assessments generally rely on large numbers of trials and participants and don’t point to specific underlying muscle interactions as provided here.

Together, through two example applications, we have demonstrated attributes of the proposed methodology that provide novel capabilities to current muscle synergy analysis. In the following, we sequentially present in more detail the three types of muscular interactions in the spatiotemporal domain across three datasets and then show the robustness of the approach and its outputs.

### Representations of motor behaviour in muscle couplings

To begin, we derived pairs of muscle activation vectors [*m*_*x*_, *m*_*y*_] from benchmark datasets of human participants executing naturalistic movements, namely arm reaching (dataset 1), whole-body point-to-point reaching (dataset 2), and various locomotion modes (dataset 3) [25–27,33] (see Fig.4 for the experimental design of each dataset and ‘*Materials and methods’* for an outline of the experimental setups and EMG data pre-processing) (fig.3(A)). For datasets 1 and 2, we determine the MI between [*m*_*x*_, *m*_*y*_] vectors with respect to several discrete task parameters representing specific task attributes (e.g. reaching direction, speed etc.), while for dataset 3 we determined the task-relevant and -irrelevant muscles couplings in an assumption-free way by quantifying them with respect to all available kinematic, dynamic and inertial motion unit (IMU) features.

**Fig. 3:**
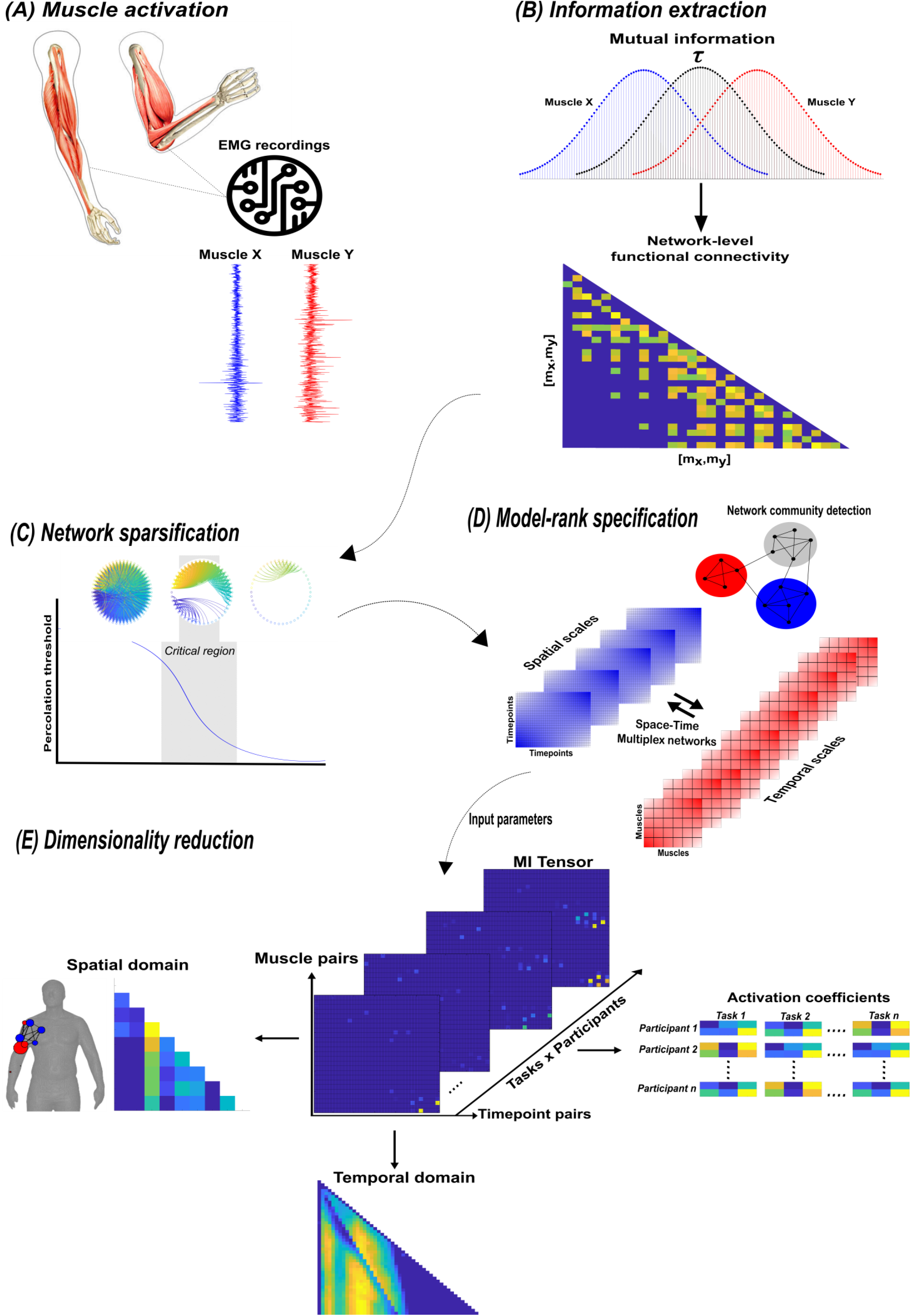
A summary of the NIF pipeline. (**A**) Large-scale datasets of EMG signals are captured while participants perform various motor tasks [25–27]. (**B**) The MI between all unique muscle-timepoint vector ([*m*_*x*_, *m*_*y*_]) combinations with respect to a corresponding task parameter (*τ*) is determined [28], forming a network of functional connectivities. (**C**) These adjacency matrices are then analysed in terms of statistical significance and modular structure using percolation theory [11]. (**D**) The optimal spatial and temporal model-ranks are determined using generalised, consensus-based network community detection methods [29–32]. (**E**) The optimal model-ranks are used as input parameters for dimensionality reduction, where space-time muscle networks along with their underlying activation coefficients are concurrently extracted [25].

Having extracted the muscle pair-task interdependencies representing a specific intersection in fig.1.1(b), we next sought to find a parsimonious representation of motor behaviour that is consistent across tasks and participants for dataset 1-3 (fig.3(E)) [11,25]. To produce this sparse, low-dimensional representation, we undertook the following intermediary steps:

We modelled the MI values as adjacency matrices in the spatial or temporal domain and identified dependencies that were statistically significant using percolation theory (fig.3(C)) [43,44]. By assuming the muscle networks operate near a state of self-organised criticality [45], we effectively isolated dependencies that were above chance-level occurrence, thus empirically sparsifying the networks.

To empirically determine the number of components to extract in a parameter-free way, we then concatenated these adjacency matrices into a multiplex network and employed network community-detection protocols to identify modules across spatial and temporal scales (fig.3(D)) [29–32,46]. Having detected the spatiotemporal modular structure, we then returned the sparsified networks to their original format and used the number of modules identified as input parameters into dimensionality reduction (fig.3(E)) [25].

By optimising a modularity-maximising cost-function [47,48], the community detection protocols we employed consistently identified three spatial (S1-S3) and two temporal (T1-T2) modules as representative of the underlying task-redundant, -synergistic and -irrelevant informational dynamics. Following their extraction, we further analysed the spatial networks from each dataset in terms of their submodular structure by applying network-theoretic tools [29,32,39]. In doing so, we identified subnetworks within each spatial network and interesting patterns of network centrality, i.e. the relative importance of a node in a network. The spatial and temporal networks of each dataset output are illustrated in panels A and B (fig.7-12) of the following sections. They are accompanied by human body models where node colour and size indicate subnetwork community affiliation and network centrality respectively [38]. The networks we extracted operate in parallel within spatial and temporal domains while having an all-to-all correspondence across domains, i.e. any spatial component can be combined with any temporal component via a task-specific coefficient (illustrated in panel C for dataset 1 and 2 outputs and in the supplementary materials (fig.4-6) for dataset 3) [25]. Unlike similar muscle synergy extraction approaches, dimensionality reduction in the NIF pipeline doesn’t seek to approximate the variance of recorded EMG data but to identify sets of muscles that share the same type of interaction. Thus, the multiplexing coefficients extracted in this framework are instead interpreted as the participant- and task-specific scaling of information overlap.

**Fig. 4:**
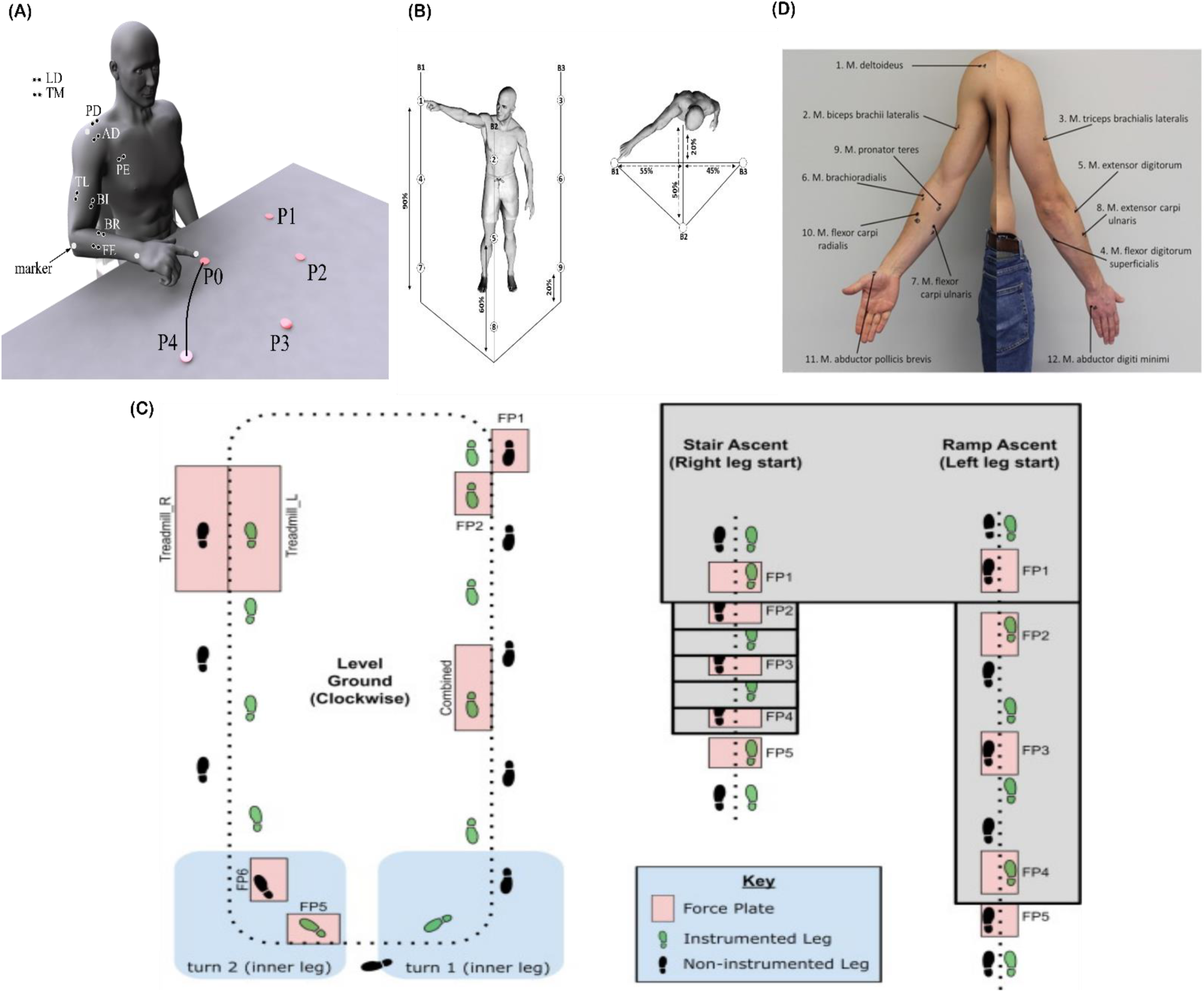
Graphical illustrations of each of the datasets analysed in the current study. **(A)** Dataset 1 consisted of participants executing table-top point-to-point reaching movements (40cm distance from starting point P0) across four targets in forward (P1-P4) and backwards (P5-P8) directions at both fast and slow speeds (40 repetitions per task) [25]. The muscles recorded included the finger extensors (FE), brachioradialis (BR), biceps brachii (BI), medial-triceps (TM), lateral-triceps (TL), anterior deltoid (AD), posterior deltoid (PD), pectoralis major (PE), latissimus dorsi (LD) of the right, reaching arm. **(B)** For dataset 2, the activity of 30 muscles was recorded while participants performed whole-body point-to-point reaching movements across three different heights and bars and in various directions, accumulating to 72 unique reaching tasks [26]. **(C)** The circuit navigated by participants in dataset 3 as they executed various locomotion modes is illustrated, of which level-ground walking, stair- and ramp-ascent/descent were analysed in the current study [27]. Several sub-conditions were undertaken by participants for each locomotion mode including different walking-speeds, clockwise vs. counter-clockwise direction, different stair heights and ramp inclines etc. Participants executed these tasks while the EMG of 11 muscles on the right leg ((Gluteus medius (GlutM), right external oblique (Obl), semitendinosus (ST), gracilis (GR), biceps femoris (BF), rectus femoris (RF), vastus lateralis (VL), vastus medialis (VM), soleus (SO), tibialis anterior (TA), gastrocnemius medialis (GM)) along with kinematic, dynamic and IMU signals were captured. (**D**) The EMG placement for dataset 4 [Deltoideus pars clavicularis (DC), Biceps brachii (BB), Triceps brachii (TB), Flexor digitorum superficialis (FDS), Extensor digitorum (ED), Brachioradialis (BR), Flexor carpi ulnaris (FCU), Extensor carpi ulnaris (ECU), Pronator teres (PT), Flexor carpi radialis (FCR), Abductor pollicus brevis (APB), Abductor digiti minimi (ADM)] [33]. A single-trial was taken from 25 healthy and 20 post-stroke participants performing a unilateral pointing movement with the index finger and arm outstretched (task 9 of the Softpro protocol (MHH)).

**Fig. 5:**
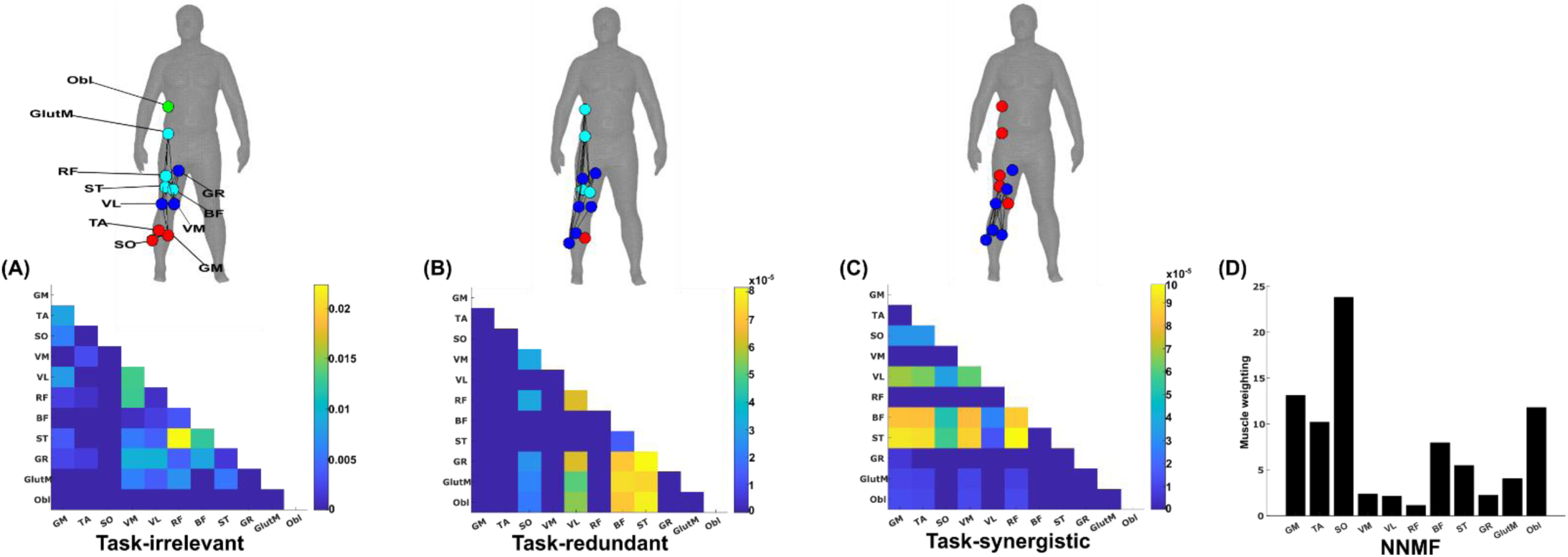
A simplified example output from the proposed framework applied to a single trial of turning gait from Dataset 3. (**A**) Task-irrelevant, (**B**) Task-redundant and (**C**) Task-synergistic synchronous muscle couplings were quantified (the unit of shared information is 1 bit) with respect to the heel kinematic marker (anterior-posterior direction). Human body models accompanying each spatial network illustrate their respective submodular structure with node colour and size and edge width indicating community affiliation [29], network centrality and connection strength respectively [38,39]. (**D**) A corresponding synergy representation from a single trial of turning gait from dataset 3 extracted using the spatial model from current approaches [37]. Each bar represents the relative weighting of each muscle in the synergy component.

**Fig. 6:**
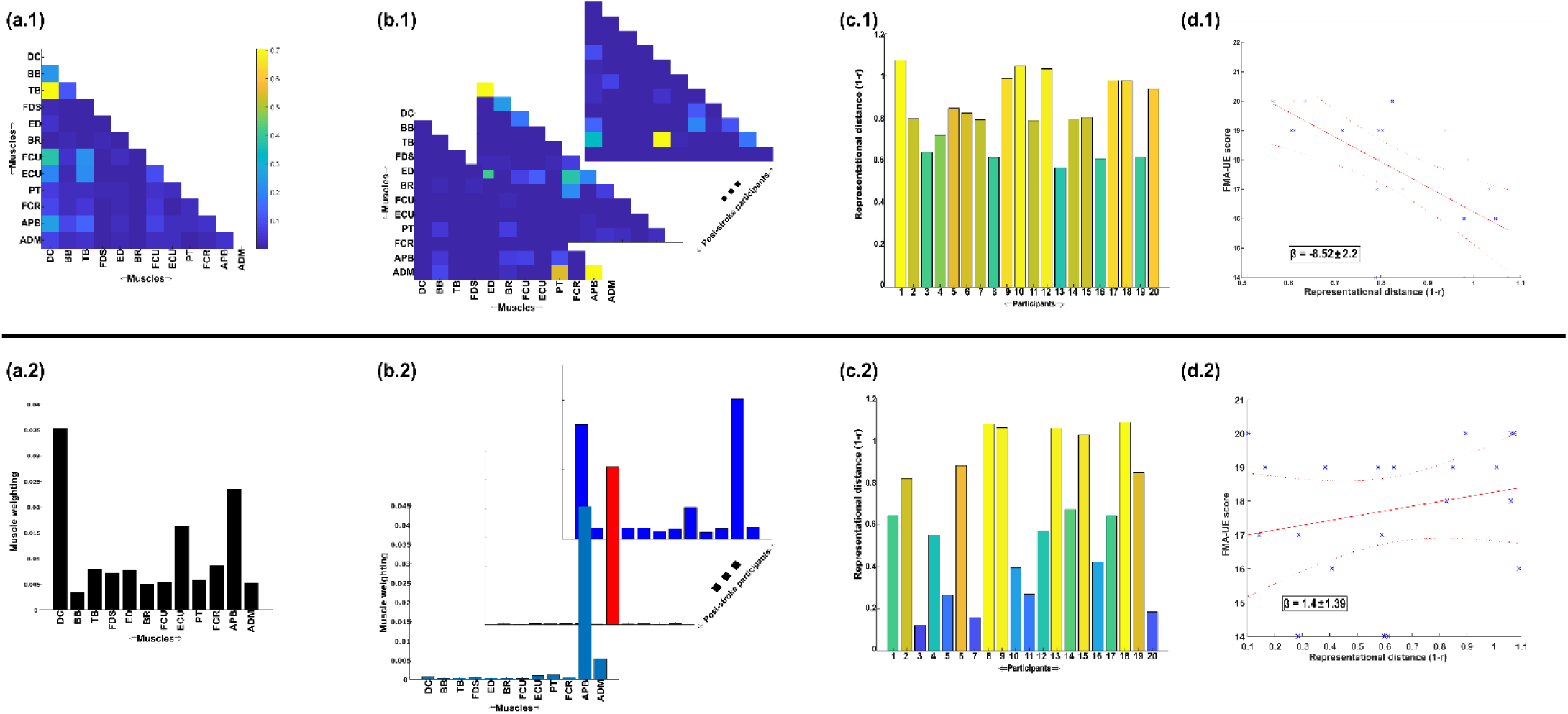
A simple demonstration of the physiological relevance of the proposed approach (**a.1-d-1**) and the traditional, NMF-based approach (**a.2-d.2**). From dataset 4 [33], we took the EMG signals and WRBA kinematic from 20 post-stroke and 25 healthy participants. We extracted a single normative reference of healthy controls task-redundant muscle couplings with respect to WRBA (**a.1**) and a corresponding normative reference using NMF only (**a.2**). We then extracted a single component from each post-stroke participant and compared them individually with the corresponding normative reference, computing distance values (1-r) (**b-c**). We finally determined the predictive relationship of these distance values with a measure of upper-extremity motor impairment derived from the Fugl-meyer assessment (FMA-UE) (**d**).

#### Task-irrelevant muscle couplings

To quantify the task-irrelevant contributions of muscular interactions to motor behaviour, we conditioned the MI between [*m*_*x*_, *m*_*y*_] with respect to *τ* (see ‘*Materials and methods’* section) [28]. This conditioning effectively removes the task-relevant information, leaving information produced by pairwise muscle variations that are task-indiscriminative. Following a run through the NIF pipeline (fig.3) [11], the output from dataset 2 and 3 are presented in fig.7-8 respectively while the output from dataset 1 is presented in the supplementary materials (fig.1). The task-irrelevant space-time muscle networks we extracted from dataset 1 and 2 shared several structural features with their task-agnostic counterparts extracted in the preliminary study [11], supporting recent work showing that functional muscle network structure are heavily influenced by task-irrelevant factors such as anatomical constraints [49]. The functional connectivities identified here captured known contributions of spatiotemporal muscular interactions to aspects of motor behaviour common across tasks and participants which we outline briefly here.

**Fig. 7:**
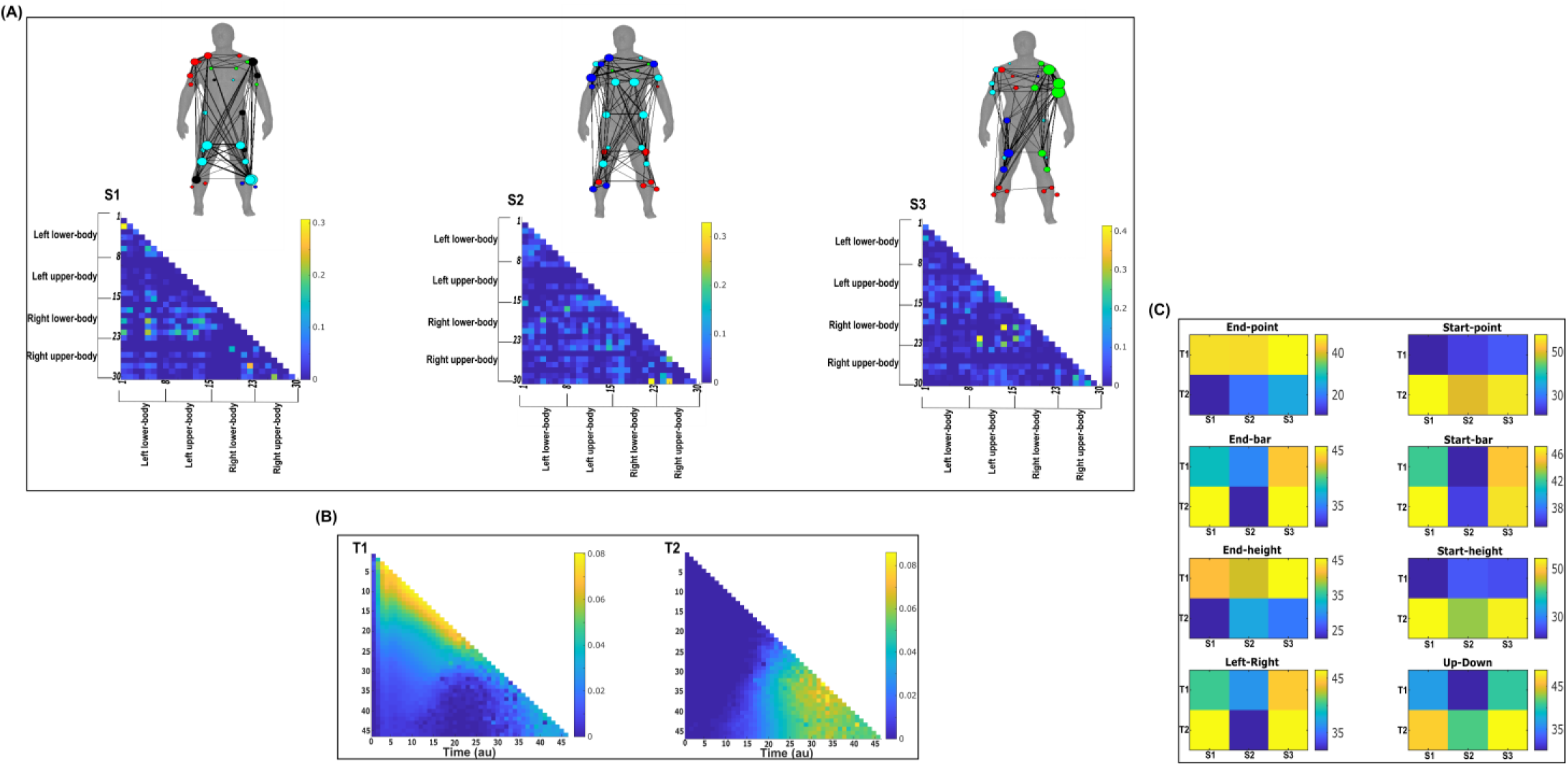
Three spatial (**S1**-**S3**) and two temporal task-irrelevant muscle networks (**T1**-**T2**) were empirically identified and extracted across participants and task parameters from dataset 2 using the NIF pipeline (Panel **A**-**B**) [11,26]. (Panel **C**) Activation coefficients are presented to the right of the networks, indicating their task parameter-specific scaling averaged across participants. Human body models accompanying each spatial network illustrate their respective submodular structure with node colour and size and edge width indicating community affiliation [29], network centrality and connection strength respectively [38,39].

**Fig. 8:**
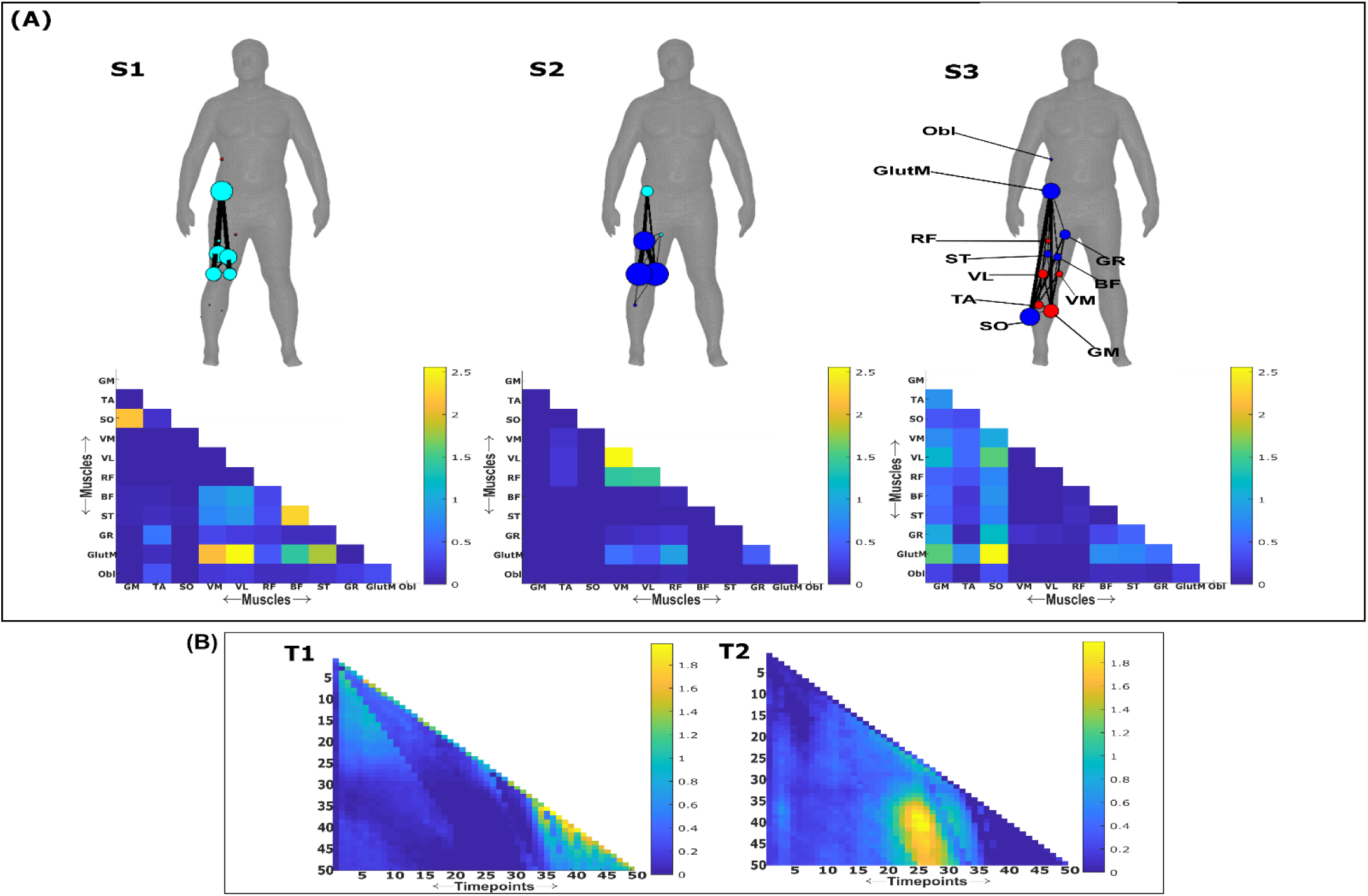
Three spatial (**S1**-**S3**) and two temporal task-irrelevant muscle networks (**T1**-**T2**) were empirically identified and extracted across participants and task parameters from dataset 3 using the NIF pipeline (Panel **A**-**B**) [11,27]. Activation coefficients are presented in supplementary materials (fig.4), indicating their task parameter-specific scaling averaged across participants in the dynamic, IMU and kinematic spaces. Human body models accompanying each spatial network illustrate their respective submodular structure with node colour and size and edge width indicating community affiliation [29], network centrality and connection strength respectively [38,39].

The temporal networks from dataset 1 and 2 captured mostly co-activations from movement onset – mid-movement and movement cessation, indicating that some co-contraction mechanisms were consistently task-irrelevant across trials. The temporal networks for dataset 2 were more diffuse compared to dataset 1, probably reflecting the more variable role of passive forces in generating movements to different heights captured in this dataset’s experimental design [26,50]. Furthermore, this co-contraction mechanism was more parsimoniously represented as a single network in dataset 3 (T1 in fig.8B), where passive forces in contrast likely played a consistently resistive role during locomotion. Interestingly, T1 for dataset 3 corresponded equivalently high for all three task spaces when corresponding with S2 which consisted of upper-leg extensors (see Supplementary materials (fig.4)). Muscle couplings indicative of agonist-antagonist pairings were also identified as separate subnetworks in S3 of dataset 3 (fig.8A). More specifically, their functional segregation appeared to be based on their distinct functional roles in forward propulsion (red nodes) and deceleration (blue nodes) during the mid-stance phase of gait, as indicated by the prominent correspondence with T2 across task spaces (see Supplementary materials (fig.4)). This finding reflects the consistent agonistic and antagonistic contributions of muscular interactions across locomotive tasks.

The gross motor function of muscle couplings was another characteristic of task-irrelevant muscle couplings that pervaded across the datasets analysed here. For instance, AD had a central role in S2 of dataset 1 while also displaying a unique pattern of connectivity with tibial musculature in S3 of dataset 2. Similarly, GlutM had a central role in S1 of dataset 3 (fig.8A). We further found a common pattern of task-irrelevant connectivity in S2 across datasets, namely the musculature about a hinge joint (elbow in dataset 1 and 2, knee in dataset 3) coupled with proximal shoulder or hip musculature, indicative of their biomechanical affordances. Finally, the passive, left-arm was connected with the tibial musculature of S3 in dataset 2 (green nodes (Fig.7(A)). To probe the underlying function of this connectivity in the left-arm, we inspected the original EMG signals. We observed periodic, tonic activations across tasks, reflective of reciprocal inhibition of contralateral limb musculature that enables unilateral movement [51].

#### Task-redundant muscle couplings

To characterise the functional role of task-relevant muscle couplings, we employed a higher-order information-theoretic concept known as co-information (co-I) [28,52,53]. This metric quantifies the MI between three random variables and may take on positive values (net synergistic) and negative values (net redundant) (see Fig.13 of ‘*Materials and methods’* section). Co-I quantifies the task-relevant information shared between [*m*_*x*_, *m*_*y*_] independently of the information generated by task-irrelevant muscular interactions. In doing so, it also defines the functional relationship between [*m*_*x*_, *m*_*y*_] overall as redundant or complementary. Following the quantification of co-I for all [*m*_*x*_, *m*_*y*_] and corresponding *τ* (pink area in orange-and-pink intersection (fig.1.3(B)), we parsed the negative values indicating redundancy into a separate matrix and rectified them. In fig.9-10, we illustrate the output following the extraction of task-redundant space-time muscle networks from datasets 2-3 across tasks and participants respectively while the output for dataset 1 is presented in supplementary materials fig.2. In the co-I formulation, task-redundant muscle couplings can be interpreted as muscle couplings that overall shared a common task-relevant functional role. For example, with reference to Figure 9 here, muscles in the networks presented in S1 (fig.9A) carry redundant information about the movement endpoint (fig.9C) with the temporal profile T1 (fig.9B) whereas S3 (fig.9A) contains muscle networks carrying redundant information about the starting point (fig.9C) with the same temporal profile T1 (fig.9B).

**Fig. 9:**
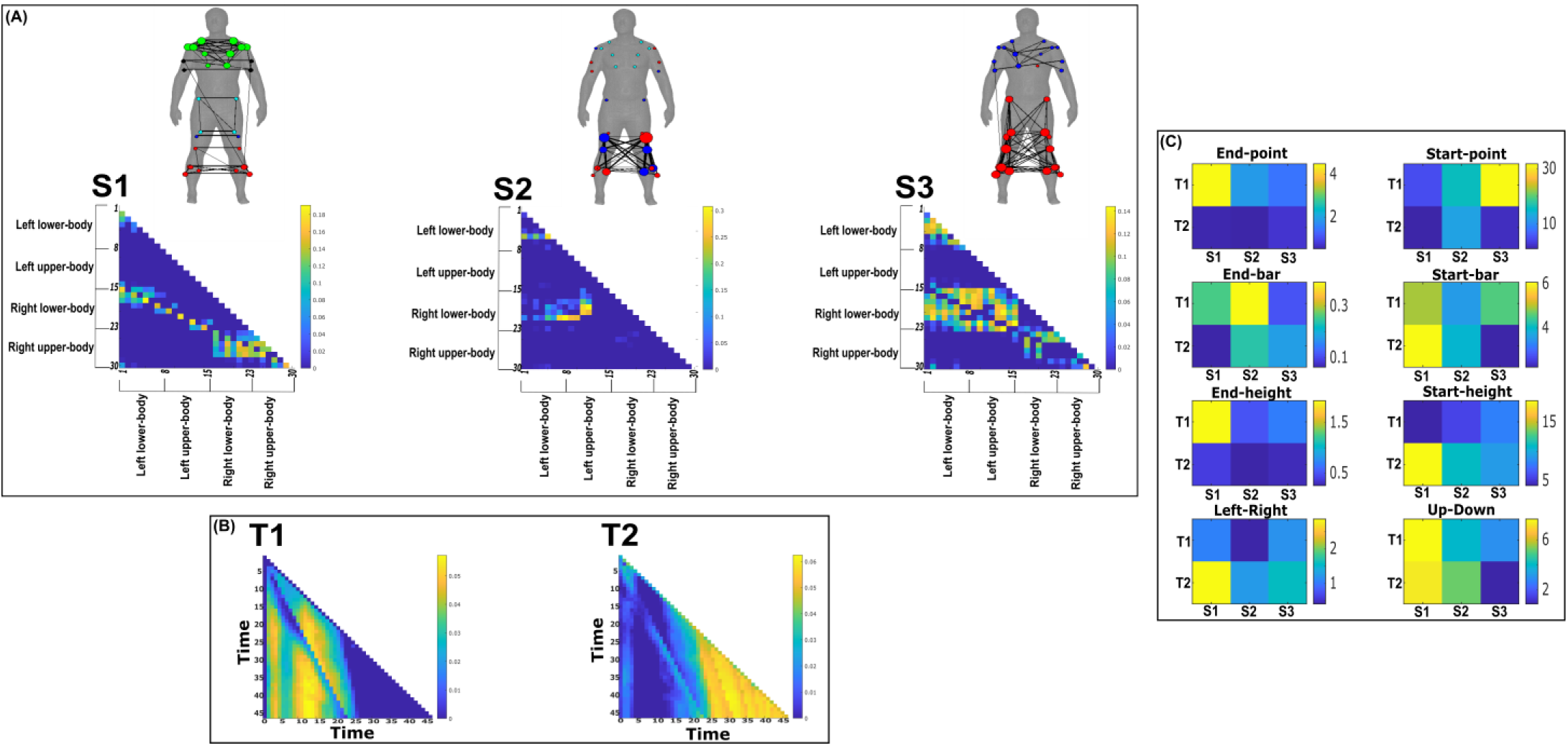
Three spatial (**S1**-**S3**) and two temporal task-redundant muscle networks (**T1**-**T2**) were empirically identified and extracted across participants and task parameters from dataset 2 using the NIF pipeline (Panel **A**-**B**) [11,26]. (Panel **C**) Activation coefficients are presented to the right of the networks, indicating their task parameter-specific scaling averaged across participants. Human body models accompanying each spatial network illustrate their respective submodular structure with node colour and size and edge width indicating community affiliation [29], network centrality and connection strength respectively [38,39].

Both dataset 1 (supplementary materials fig.2) and dataset 3 (fig.10) outputs display similar patterns of muscle couplings at the same spatial scale of an individual, task-relevant limbs’ musculature, with an emphasis on the coupling of specific muscles with all other muscles. For dataset 1, FE, BI and BR displayed this integrative pattern across S1-S3 respectively while BF, TA and ST demonstrated this pattern in dataset 3 also. The muscle networks encapsulated several functionally interpretable couplings such as the agonist-antagonist pairings of the BI and TM of S2 (dataset 1 (supplementary materials fig.2(A)) and the task redundancy of ankle dorsi-flexors, and knee/hip flexors during sloped walking for example in S2 of dataset 3 (fig.10(A)) [54]. The functional interpretation of these muscle connectivity patterns was in line with the extracted task-specific activations. For instance, S2 of dataset 1 was modulated most prominently by reaching direction when corresponding with T2, commensurate with the biomechanical affordances of this upper-arm muscle network. Furthermore, S2 of dataset 3 was specifically modulated by the right-thigh kinematic marker along the y-axis (up-down direction) for both T1 and T2 (see Supplementary materials (fig.5)). The centrality of task-redundant muscle couplings in dataset 1 and 3 suggests particular muscle activations drive the task-specific variations in the reaching arm and stepping leg muscle activities towards a common behavioural goal. It is also worth noting that the magnitude of these functional connectivity patterns appeared to be proportional to anatomical distance, as evidenced by the magnitude of connection strengths, a finding supportive of previous related research [49].

**Fig. 10:**
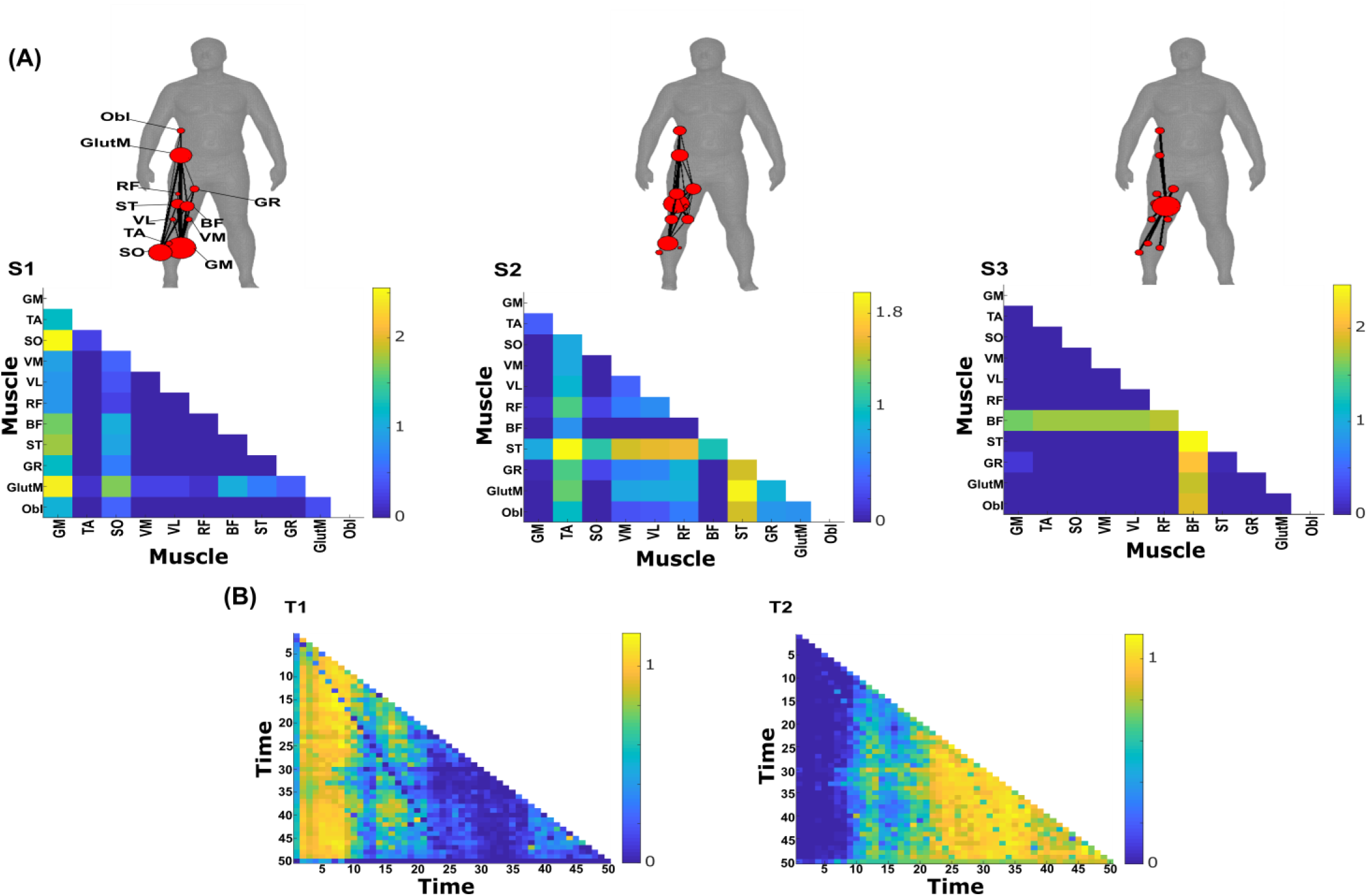
Three spatial (**S1**-**S3**) and two temporal task-redundant muscle networks (**T1**-**T2**) were empirically identified and extracted across participants and task parameters from dataset 3 using the NIF pipeline (**A**-**B**) [11,27]. Activation coefficients are presented in supplementary materials document 2 (fig.2), indicating their task parameter-specific scaling averaged across participants in the dynamic, IMU and kinematic spaces. Human body models accompanying each spatial network illustrate their respective submodular structure with node colour and size and edge width indicating community affiliation [29], network centrality and connection strength respectively [38,39].

Meanwhile at the greater spatial scale of dataset 2 (fig.9(A))), task-redundant muscle couplings were anatomically compartmentalised to the upper- and lower-body. This functional segregation was emphasised at the subnetwork level also, where the upper- and lower-body musculature of S3 for instance formed distinct submodules (blue and red nodes). Amongst the task-specific activations in dataset 2, S1 carried redundant task information about end-point target, -height and up-down direction when corresponding with T1. T2 for dataset 2 on the other hand contained mostly temporally proximal dependencies along the diagonal, suggestive of co-contraction mechanisms, which became more diffuse near movement cessation. These endpoint trajectory and co-contraction related temporal patterns were qualitatively similar to T1 and T2 of both dataset 1 and 3 respectively (see fig.10 and Supplementary materials fig.2 respectively).

#### Task-synergistic muscle couplings

Similarly, we isolated task-synergistic muscle couplings by parsing, instead, the positive co-I values from the computations conducted across all [*m*_*x*_, *m*_*y*_] and corresponding *τ* into a sparse matrix where all redundant couplings were set to zero (see Fig.13 of ‘*Materials and methods’* section). Task-synergistic muscle couplings here can be interpreted as a [*m*_*x*_, *m*_*y*_] pair that provide complementary (i.e. functionally dissimilar) task information, thus more information is gained by observing [*m*_*x*_, *m*_*y*_] together rather than separately (orange area in orange-and-pink intersection (fig.1.3(B)). In Fig.11-12, we illustrate the task-synergistic space-time muscle networks from datasets 2-3 respectively (dataset 1 output is presented in the supplementary materials fig.3).

**Fig. 11:**
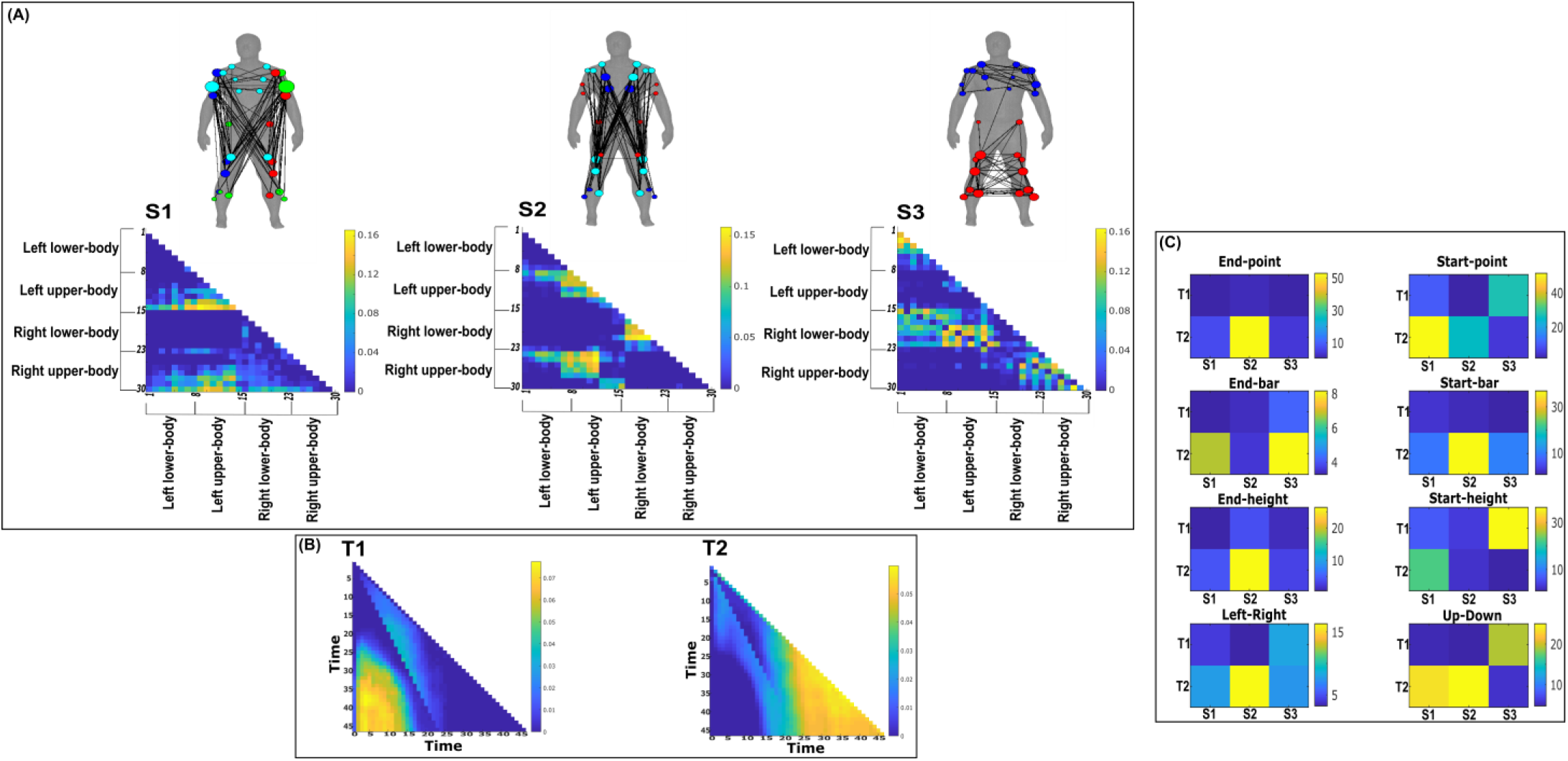
Three spatial (**S1**-**S3**) and two temporal task-synergistic muscle networks (**T1**-**T2**) were empirically identified and extracted across participants and task parameters from dataset 2 using the NIF pipeline (Panel **A**-**B**) [11,26]. (Panel **C**) Activation coefficients are presented to the right of the networks, indicating their task parameter-specific scaling averaged across participants. Human body models accompanying each spatial network illustrate their respective submodular structure with node colour and size and edge width indicating community affiliation [29], network centrality and connection strength respectively [38,39].

**Fig. 12:**
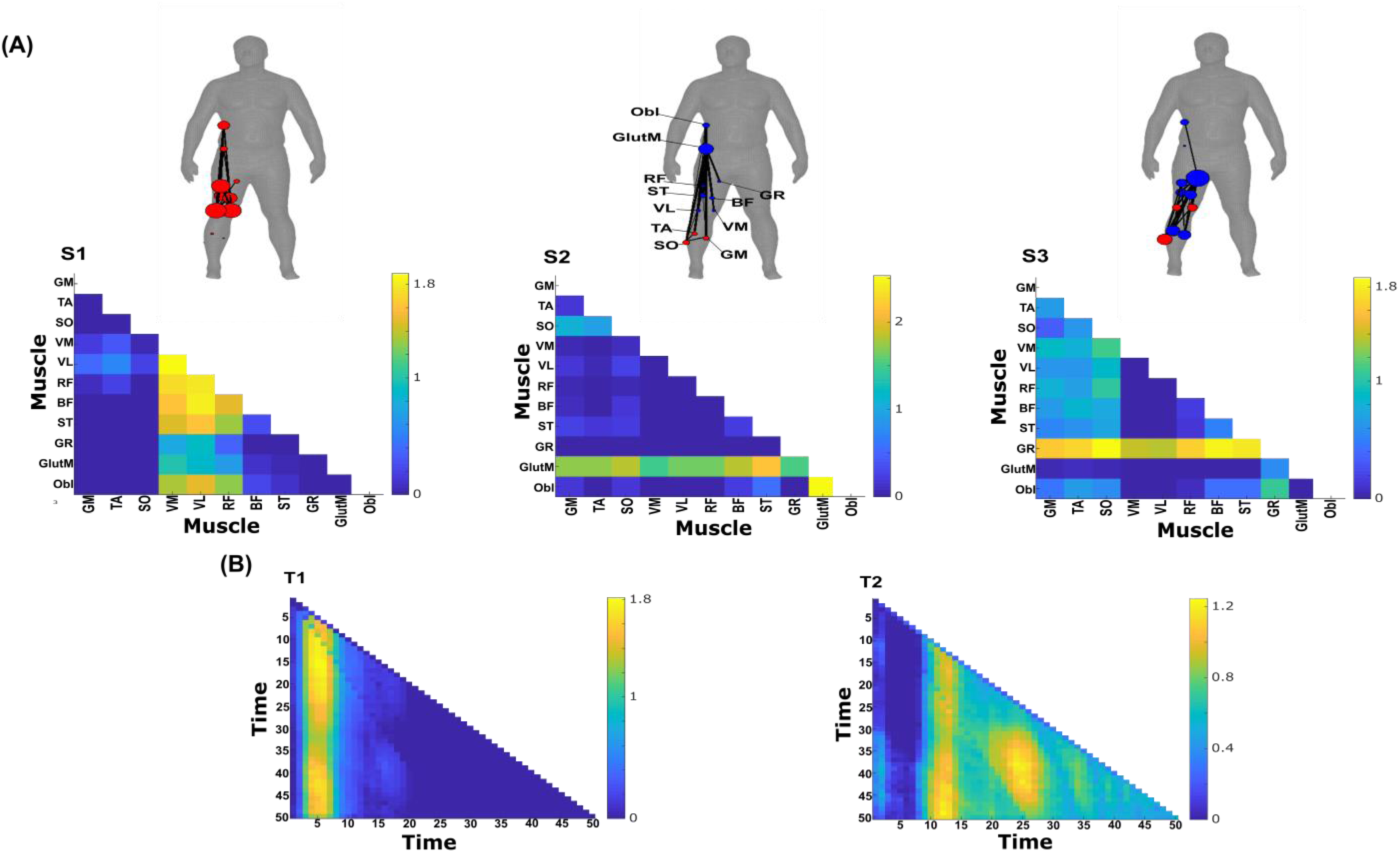
Three spatial (**S1**-**S3**) and two temporal task-synergistic muscle networks (**T1**-**T2**) were empirically identified and extracted across participants and task parameters from dataset 3 using the NIF pipeline (Panel **A**-**B**) [11,27]. Activation coefficients are presented in supplementary materials (fig.6), indicating their task parameter-specific scaling averaged across participants in the dynamic, IMU and kinematic spaces. Human body models accompanying each spatial network illustrate their respective submodular structure with node colour and size and edge width indicating community affiliation [29], network centrality and connection strength respectively [38,39].

**Fig. 13:**
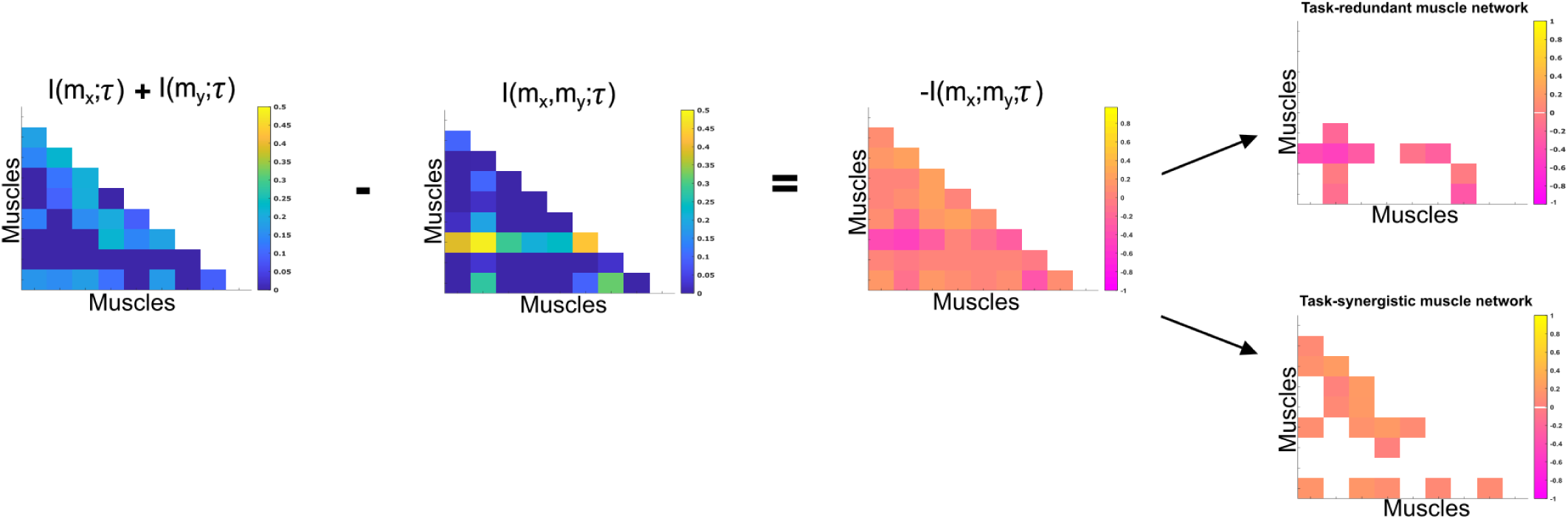
Co-I determines the difference between the sum total information shared with *τ* in *m*_*x*_ and *m*_*y*_ when observed separately and the information shared with *τ* when they are observed together. The adjacency matrices show how this calculation is carried out for all unique [*m*_*x*_, *m*_*y*_] combinations. Redundant and synergistic muscle couplings are then separated into two equivalently sized networks. The accompanying colour bars indicate the values present in the adjacency matrix.

Across datasets, muscle networks could be characterised by the transmission of complementary task information between functionally specialised muscle groups, many of which were identified among the task-redundant representations (Fig.9-10 and Supp. Fig.2). The most obvious example of this is the S3 synergist muscle network of dataset 2 (Fig.11), which captures the complementary interaction between task-redundant submodules identified previously (S3 (Fig.9)). A particularly consistent structural feature was the emphasis of an individual muscles’ connectivity with all other muscles which was evident among synergistic couplings in dataset 3 (see VM, VL and RF of S1, GR, TA and SO of S2 and GlutM of S3 in fig.12A). This structural similarity demonstrates that parallel and synchronous exchanges of redundant and synergistic task information underlie task-specific variations across trials (e.g. S3 of dataset 2 in fig. 9A and fig.11A). Interestingly, despite the similarity between the redundant and synergistic muscle networks, the way they are combined to encode task information differs depending on the type of interaction (synergistic vs. redundant, e.g. panel C of fig.11 in comparison to panel C of fig.9).

Concerning the temporal activations of these networks, the task-synergistic structure of dataset 2 (fig.11B) was also relatively unchanged compared to the task-redundant structure (fig.9B). This suggests that the task end-goal and co-contraction related mechanisms provided both redundant and synergistic task information concurrently during whole-body reaching movements. In contrast, a different view of synergistic information exchange is provided in datasets 1 and 3 (supplementary fig.3 and fig.12 here respectively), where T1 and T2 consist of more idiosyncratic activations that together appear to reflect the task end-goal related patterns found elsewhere. More specifically, in both datasets we found two distinct patterns of task- end goal related activity where early and late timepoints during movement initiation operated in parallel to provide complementary task information (see fig. 12B and supp. fig. 3B).

### Generalisability of the extracted space-time muscle networks

To ascertain the generalisability of the extracted representations presented here beyond any subset of the input data, we conducted a similarity analysis through a leave-n-out cross validation procedure. In more detail, we compared the space-time networks extracted from the full dataset (illustrated in fig.4-9 and supplementary materials here) to the networks extracted from a subset of the data (see ‘*Materials and methods’* section). Across datasets, a high level of concordance was found on average (∼0.9 correlation, see table1). This trend was evident across all datasets and for task-redundant, -synergistic and -irrelevant spatial and temporal networks. Dataset 1 and 2 findings demonstrate that the extracted networks are generalisable beyond any individual participant or task. Dataset 3 results go further by demonstrating that the extracted patterns are generalizable beyond any randomly selected and randomly sized subset of the input data. The highest correlations on average were consistently among temporal networks, replicating previous findings [11,55,56]. Although the spatial networks demonstrated a lower average correlation, this was substantially higher compared to previous applications of datasets 1 and 2 [11,25,55], suggesting that the inclusion of task parameters here captures the inter-participant differences more effectively. When comparing representations extracted from each continuous task space in dataset 3, kinematic features consistently had the highest average correlation and lowest variability compared to dynamic and IMU feature spaces. These findings support our change in the interpretation of the extracted activation coefficients away from conventional approaches where representational biases towards particular participants and/or tasks are inferred.

**Table. 1:**
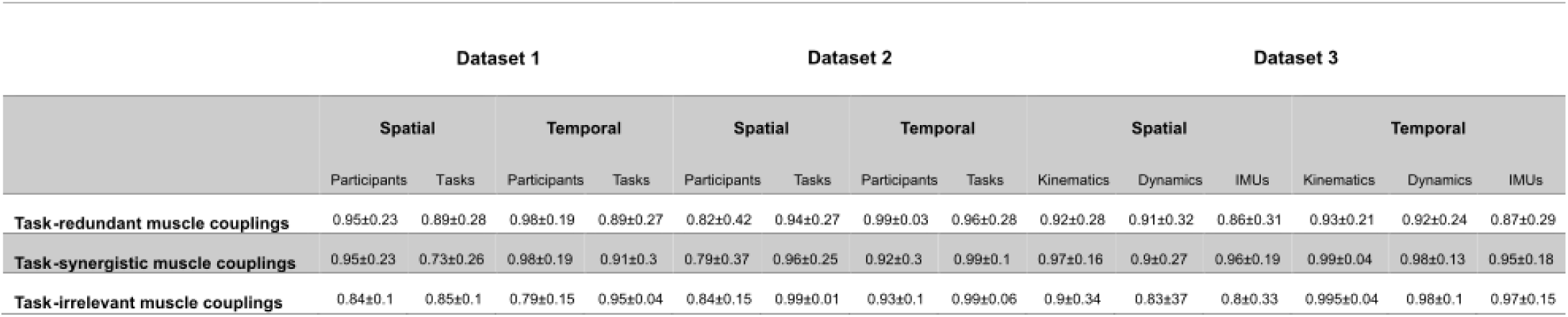
A summary table illustrating the findings from an examination of the generalisability of the muscle networks extracted from each dataset. The spatial and temporal representations extracted from the full input data in each muscle-task information subspace were compared using Pearson’s correlation against functionally similar representations extracted from a subset of the input data

|  | Dataset 1 |  |  |  | Dataset 2 |  |  |  | Dataset 3 |  |  |  |  |  |
| --- | --- | --- | --- | --- | --- | --- | --- | --- | --- | --- | --- | --- | --- | --- |
|  | Spatial |  | Temporal |  | Spatial |  | Temporal |  | Spatial |  |  | Temporal |  |  |
|  | Participants | Tasks | Participants | Tasks | Participants | Tasks | Participants | Tasks | Kinematics | Dynamics | IMUs | Kinematics | Dynamics | IMUs |
| **Task-redundant muscle couplings** | 0.95±0.23 | 0.89±0.28 | 0.98±0.19 | 0.89±0.27 | 0.82±0.42 | 0.94±0.27 | 0.99±0.03 | 0.96±0.28 | 0.92±0.28 | 0.91±0.32 | 0.86±0.31 | 0.93±0.21 | 0.92±0.24 | 0.87±0.29 |
| **Task-synergistic muscle couplings** | 0.95±0.23 | 0.73±0.26 | 0.98±0.19 | 0.91±0.3 | 0.79±0.37 | 0.96±0.25 | 0.92±0.3 | 0.99±0.1 | 0.97±0.16 | 0.9±0.27 | 0.96±0.19 | 0.99±0.04 | 0.98±0.13 | 0.95±0.18 |
| **Task-irrelevant muscle couplings** | 0.84±0.1 | 0.85±0.1 | 0.79±0.15 | 0.95±0.04 | 0.84±0.15 | 0.99±0.01 | 0.93±0.1 | 0.99±0.06 | 0.9±0.34 | 0.83±0.37 | 0.8±0.33 | 0.995±0.04 | 0.98±0.1 | 0.97±0.15 |

To further probe how the underlying assumption of an all-to-all correspondence between spatial and temporal representations made by sNM3F influenced the generalisability of the extracted networks, we compared its performance to non-negative Canonical-Polyadic (CP) tensor decomposition, which assumes an opposing one-to-one correspondence between components. We found that although CP has demonstrated a considerable capacity to de-mix neural data into simplified and interpretable low-dimensional components [57], its application here resulted in poor generalisability of the extracted patterns (∼0.5 correlation on average). This finding suggests that the all-to-all correspondence implemented by sNM3F identifies a more generalizable representation and should be favourably considered in future applications of this framework.

## Discussion

The aim of the current study was to dissect the muscle synergy concept and offer a novel, more nuanced perspective on how muscles ‘*work together’* to achieve a common behavioural goal. To do so, we introduced a computational approach based on the NIF pipeline (fig.3), enabling the effective decomposition of muscle interactions and the comprehensive description of their functional roles. Through the direct inclusion of task parameters in the extraction of muscle synergies, a novel perspective was produced where muscles ‘*work together’* not just towards a common task-goal, but also concomitantly towards complementary task objectives and lower-level functions irrelevant for overt task performance. The functional architectures we uncovered were comprised of distributed subnetworks of synchronous muscle couplings and driven by distinct temporal patterns operating in parallel. Example applications to simple real and simulated EMG datasets revealed the additional capabilities provided by the proposed framework to current muscle synergy analysis in terms of functional and physiological relevance and interpretability. When applied to large-scale data, the proposed methodology extracted representations scale-invariant to dataset complexity and motor behaviours whilst being generalizable beyond any subset of the data. We thus present this framework as a useful analytical approach for mechanistic investigations on large-scale neural data through this novel perspective to the muscle synergy.

The ‘*muscle synergy’* is a major guiding concept in the motor control field [3–7,15], providing a conceptual framework to the problem of motor redundancy that centres around the general notion of ‘*working together’*. In its current conceptualisation, ‘*working together’* describes how the nervous system functionally groups muscles in a task-dependent manner through common neural drives to simplify movement control. This idea has undergone continued refinement since its early conception [3,6,7], with a notable progression being the introduction of the qualifying attributes: a sharing pattern, reciprocity and task dependence [7]. Nonetheless, recent influential works revealing other important mechanisms for the simplification of motor control highlight the generality of the current perspective offered by this concept [16–24]. We thus sought to provide greater nuance to the notion of ‘*working together’* by defining motor redundancy and synergy in information-theoretic terms [6,58]. In our framework, redundancy and synergy are terms describing functionally similar and complementary motor signals respectively, introducing a new perspective that is conceptually distinct from the traditional view of muscle synergies as a solution to the motor redundancy problem [3,6,7]. In this new definition of muscle interactions in the task space, a group of muscles can ‘*work together’* either synergistically or redundantly towards the same task. In doing so, the perspective instantiated by our approach provides novel coverage to the partitioning of task-relevant and - irrelevant variability implemented by the motor system along with an improved specificity regarding the functional roles of muscle couplings [20–22]. Our framework emphasises not only the role of functionally redundant muscle couplings that result from the underlying degeneracy of the motor system, but also of complementary, synergistic dependencies that are important for communication and integration across specialised neural circuitry [59,60]. Thus, the present study aligns the muscle synergy concept with the current mechanistic understanding of the nervous system whilst offering an analytical approach amenable to the continued advances in large-scale data capture [14,61].

Among current approaches to muscle synergy analysis, the established synchronous, temporal and time-varying muscle synergy models are understood to each characterise unique motor features [62]. More specifically, the synchronous model captures agonist-antagonist muscle pairings, the temporal model decomposes EMG signals into functionally distinct temporal phases and the task-specific modulation of spatiotemporal invariants are quantified in the time-varying model. In a unifying framework, here we quantify space-time muscle networks that concurrently capture many of these salient motor features in a holistic and principled way whilst mapping their functional consequences to motor behaviour. These salient features included, among others, agonist-antagonist pairings and functionally meaningful inter-limb couplings that consistently appeared across task-redundant, -synergistic and -irrelevant spaces. Thus, in dissecting the muscle interactions governing coordinated movement, our framework revealed their parallel and synchronous processing of functionally similar, complementary, and task-irrelevant information. This insight aligns with several recent works also demonstrating this distributed neural architecture of parallel information processing units [1,59,60].

Our framework also revealed novel characteristics of the motor system. For instance, the task-redundant and -synergistic networks we extracted appeared to be structured around the coupling between a prime-mover muscle and several supporting muscles, supporting recent work showing the nonhomogeneous sharing of neural drives within modules [63]. These novel spatial characteristics were driven by parallel temporal patterns representing endpoint trajectory and co-contraction related mechanisms, an insight supportive of recent work showing their parallel innervation [16,50,64]. Together, these representations encapsulated the functional interplay between task end-goal requirements and biomechanical affordances, a dynamic frequently highlighted in object manipulation experiments [65]. In other words, the task-relevant networks reflected how muscles ‘work together’ both redundantly and synergistically towards a desired end-goal state whilst, in parallel, continually controlling the present trajectory of the system. Meanwhile, the task-irrelevant networks demonstrated that these muscle couplings also work concomitantly towards lower-level objectives assigned mostly during the transition to this desired end state. Although distinguishing task-irrelevant muscle couplings may capture artifacts such as EMG crosstalk, our results convey several physiological objectives of muscles including gross motor functions [66], the maintenance of internal joint mechanics and reciprocal inhibition of contralateral limbs [19,51]. Thus, task-irrelevant muscle interactions reflect both biomechanical- and task-level constraints that provide a structural foundation for task-specific couplings. The separate quantification of these muscle interaction types opens up novel opportunities in the practical application of muscle synergy analysis, as demonstrated in the current study through the identification of a significant predictor of motor impairment post-stroke from single-trials [5,12,67]. For instance, these distinct representations may encapsulate different neural substrates that can be specifically assessed at the muscle-level for the purpose of bodily restoration and augmentation [68]. Uncovering their neural underpinnings is an interesting topic for future research.

Indeed, in future work, we aim to complement this study’s combinatorial perspective to the muscle synergy by dissecting the unique contribution of individual muscles to motor behaviour and how they may work independently towards task performance (see magenta and cyan intersections in fig.1.3(B)). More broadly, our work here parallels related information-theoretic approaches to decomposing task-relevant brain activity [53,69], whilst addressing a current research gap across the neurosciences in effective methods to mapping large-scale, spatiotemporal neural activity to behaviour [14,61]. Future applications of this framework should include large-scale, multi-modal data captured from participants performing a wide range of natural behaviours.

In sum, this study introduced a novel perspective to the muscle synergy concept and a computational framework to extract muscle couplings that map their pairwise contributions to motor behaviour. We suggest that this approach offers novel research opportunities for investigating the underlying neural constraints on motor behaviour and the fundamental structure-function relationships generated by agent – environment interactions [15,70].

## Materials and Methods

### Quantifying muscle couplings in the task space

To quantify muscle couplings we used MI, a non-linear measure of dependence that captures any type of relationship between random variables. Here to estimate MI, we used a Gaussian copula-based approximation [28]. This semi-parametric estimator exploits the equivalence between MI and the negative entropy of the empirical copula (*c*), a function that maps a multivariate set (e.g. [*m*_*x*_, *m*_*y*_] representing activities of muscles X and Y) to their joint distribution (Equation 1.1) [28].

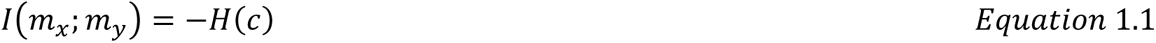

Thus, to determine task-irrelevant muscle couplings (*I*(*m*_*x*_; *m*_*y*_|*τ*)), we conditioned the negative entropy of the empirical copula for [*m*_*x*_, *m*_*y*_] with respect to a task variable *τ* (Equation 1.2). As mentioned, [*m*_*x*_, *m*_*y*_] are continuous vectors composed of individual muscle amplitudes at specific timepoints across trials while *τ* is a corresponding discrete (e.g. movement direction for datasets 1 and 2) or continuous (e.g. movement kinematics in dataset 3) task parameter. For discrete task variables, *τ* takes one value for each trial and the MI is calculated across trials using a Gaussian mixture model [28]. In the case of continuous task variables, *τ* varies in time within a specific trial. Thus, we compute MI at each timepoint *t* using the muscle activity *m*_*x*_(*t*) and the task variable value *τ*(*t*) at this time point using a closed-form parametric estimator [28].

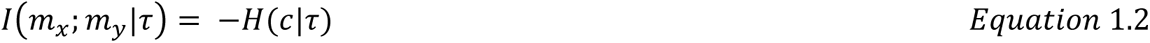

To evaluate the task-relevance of the identified muscle couplings, we used a higher-order information theoretic measure known as co-information (co-I) (Equation.1.3) (fig.11), which quantifies the relationship between three random variables, here [*m*_*x*_, *m*_*y*_], and *τ*. Co-I implements the inclusion-exclusion principle of combinatorics [52], whereby the sum of MIs between individual *m* vectors and *τ* (*I*(*m*_*x*_; *τ*) + *I*(*m*_*y*_; *τ*)) is compared against their composite MI (*I*(*m*_*x*_, *m*_*y*_; *τ*)) as follows:

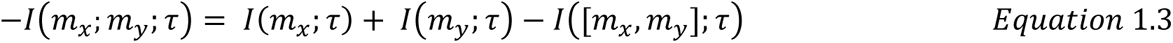

Negative *I*(*m*_*x*_; *m*_*y*_; *τ*) corresponds to a net redundant coupling between [*m*_*x*_, *m*_*y*_] about *τ* while positive *I*(*m*_*x*_; *m*_*y*_; *τ*) indicates a net synergy. To analyse these distinct couplings separately, we parsed redundant and synergistic *I*(*m*_*x*_; *m*_*y*_; *τ*) into two equivalently sized matrices and rectified the redundant couplings to make them suitable for non-negative dimensionality reduction.

Then, to produce a multiplexed view of the muscular interactions across trials, we iterated these MI computations over all unique combinations of [*m*_*x*_, *m*_*y*_] and *τ*. The resulting MI estimates collectively form *A*, a symmetric adjacency matrix (i.e. *A*^*T*^*A* = *I*) that represents the functional connectivities between all muscle activations (fig.10). When repeated across all available task variables *τ* and participants, *A* is of dimension [*No*. *of muscle pairs* x *No*. *of timepoint pairs* x [*No*. *of τ* x *No*. *of participants*]]. Thus, by applying network-theoretic statistical tools to *A*, we can identify functional modules carrying the same type of (redundant/synergistic) task information (fig.2(B)).

### Estimating statistical significance of muscle couplings

To isolate statistically significant dependencies, we applied a modified percolation analysis to each *A* [43]. This method sparsifies functional connectivities in *A* with respect to its percolation threshold (*P*_*c*_). *P*_*c*_ is a critical value that specifies the probability of a nodes’ occupation in *A* with respect to the networks size. In random networks, a ‘*giant component’* comprised of long-range connections exists above *P*_*c*_ but disappears as *P*_*c*_ tends to zero [44], while it is thought that living systems optimise adaptability by fluctuating around *P*_*c*_ in a state of self-organised criticality [45]. Preliminary testing of this method showed it to be at least equivalent to permutation-testing each MI value in the network and thus, much more computationally efficient. *P*_*c*_ was therefore iteratively specified for each layer of *A* relative to equivalently sized random networks and utilised to remove insignificant network edges up to a stopping-point where this giant component begins to become affected (fig.3(C)). This procedure was carried out for each layer of *A* separately configured as muscle-wise couplings across temporal scales (i.e. a 3-D tensor of dimension [*No*. *of muscles*] x [*No*. *of muscles*] x [*No*. *of timepoint pairings* x *No*. *of τ* x *No*. *of participants*]) and vice versa as timepoint-wise couplings across spatial scales (fig.3(D)). The separate sparsification of each individual network layer in both alternative network configurations produced discrepancies in the output, as some connections were found to be significant in only one domain. To ameliorate this discrepancy, we employed a conservative heuristic where dependencies must be significant in both space and time to be included in the final input matrix for dimensionality reduction. Thus, the sparsified input matrices to dimensionality reduction were comprised of significant spatiotemporal task-redundant, -synergistic or - irrelevant muscle couplings.

### Model-rank specification

To determine the optimal number of modules to extract, we implemented two alternative community detection algorithms generalised to multiplex networks [29,30,46]. Both forms seek to optimise a modularity criterion known as the Q-statistic that quantifies the proportion of within-community network edges compared to what would be expected from a network consisting of random connections [47]. More specifically, for a particular division of a single layer network (Equation 2.1), let *δ*(*g*_*i*_, *g*_*j*_)=1 if node *i* and *j* belong to the same group (*g*) and 0 otherwise and *A*_*ij*_ be the number of edges between node *i* and *j*. The equivalent of A_*ij*_ from a randomised network (*P*_*ij*_) is expected to be 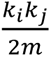 (Newman-Girvan null model) [47], where *k_i_* and *k_j_* are the node degrees and 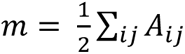. The typical output of the Q-statistic is found within the range [0,1] with 1 indicating maximum modularity [47].

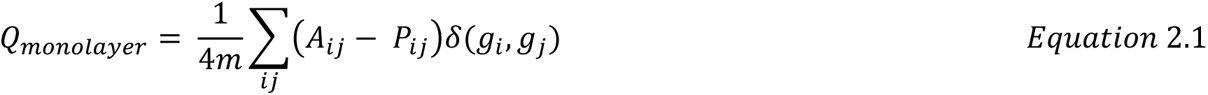

In its generalised multilayer form, the Q-statistic is given an additional term to consider couplings between layers *l* and *r* with intra- and inter-layer resolution parameters *γ* and *ω* (Equation 2.2). Here, *μ* is the total edge weight across the network and *γ* and *ω* were set to 1 in the current study for classical modularity [30], thus removing the need for any hyperparameter tuning.

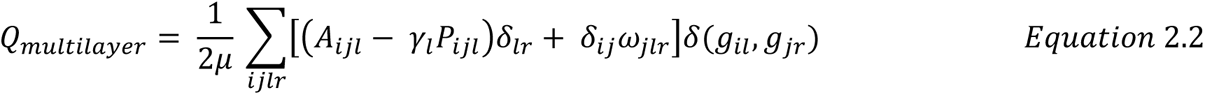

We chose to implement two complementary model-rank specification approaches to address limitations related to stochasticity and scalability present in the multilayer formulation and the inability to consider inter-layer dependencies present in the mono-layer formulation [18,48]. To apply these algorithms to our data, we grouped the set of *A* into multiplex networks configured with respect to spatial or temporal scales (fig.3(D)). We then applied these algorithms to both space-time network configurations for individual participant/tasks. This procedure generated a binary adjacency matrix from the resulting community partition vector in each case where 1 indicated the nodes belonged to the same community and 0 otherwise [46]. Following a consensus-based approach [31], we then grouped these binary adjacency matrices into a new multiplex network and re-applied the two alternative community detection algorithms to find an optimal spatial and temporal model-rank [31].

#### Extraction of low-dimensional representations

Following the specification of an optimal model-rank in the spatial and temporal domains, we used these values as input parameters into dimensionality reduction (fig.3(E)). To extract a low-dimensional representation of motor behaviour across muscle couplings, we applied a sample-based non-negative matrix tri-factorisation method (sNM3F) with additional orthogonality constraints to the matrices consisting of vectorised and concatenated *A* [25]. More specifically, we decomposed the input three-mode tensor *A* of dimension [K = No. of unique muscle pairs (*m*) x (L = No. of time-sample pairs (*t*) x No. of participants + tasks)] into a set of spatial (*V*) and temporal (*W*) factors and the participant- and task-specific weighting coefficients (*S*) reflecting the amount of information carried by each combination of spatial-temporal factors for each participant and task parameter (Equation 3.1). In equation 3.2, this factorisation is also illustrated in vector sum notation for a single participant and task variable:

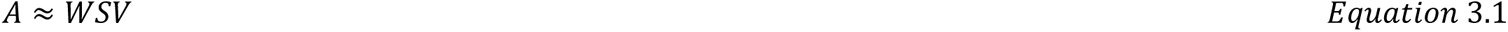

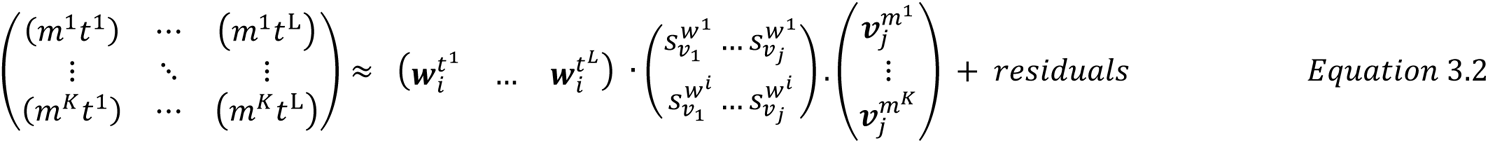

#### Examining the generalisability of extracted representations

To determine the generalisability of the extracted space-time muscle networks, we implemented a representational similarity analysis where we compared representations extracted from the full dataset 1-3 to equivalent networks extracted from a subset of the respective datasets. We computed the similarity between pairs of representations using Pearson’s correlation.

For dataset 1 and 2 (see below), we removed from the input data an individual participant or task at a time and compared the similarity of the decomposition outputs with those obtained from the full dataset. We repeated this for all participants and task variables and reported the average similarity as a measure of robustness of the decomposition.

For dataset 3 (see below), due to the greater number of participants and task parameters, we implemented a more stringent examination. More specifically, we firstly extracted representations from each task space individually and, using these representations as a reference, compared them against functionally similar outputs after removing randomly sized portions of randomly selected vectors in the input data (up to the no. of column vectors - 1). We repeated this procedure for 50 iterations, and computed summary statistics by converting the coefficients to Fisher’s Z values, computing the average and standard deviation, and then reverting these values back to correlation coefficients.

#### Subnetwork analysis

To illustrate the relative importance of individual muscles within each network, we determined the total communicability (*C*(*i*)) of individual nodes (*i*) in each network (*A*). *C*(*i*) is defined as the row-wise sum of all matrix exponentials (*e*) in the adjacency matrix (*A*) that consider the number of walks between each pair of nodes *i* and *j* (Equation 4.1) [39,71]:

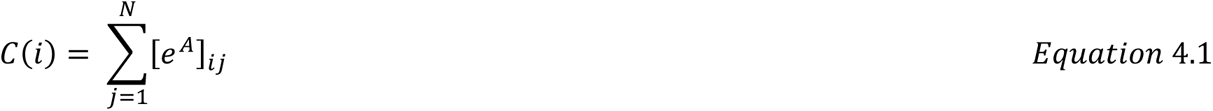

To emphasise salient functional connectivities in the spatial networks, we sparsified all dependencies with a below average network communicability and illustrated the output on the accompanying human body models [38,71]. To uncover salient subnetwork structures consisting of more closely functionally related muscle activations, we applied the monolayer community detection algorithm in equation 2.1 to the extracted spatial networks [29,32].

#### Data acquisition and processing

To illustrate our framework, we applied it to three datasets of EMG signals recorded during different motor tasks. In dataset 1 (fig.4(A)), 7 adult participants (age: 27 ± 2 years, height: 1.77 ± 0.03 m) performed table-top point-to-point reaching movements in both forward and backwards directions and at fast and slow speeds while the activity of nine muscles on the preferred right, reaching arm (finger extensors (FE), brachioradialis (BR), biceps brachii (BI), medial-triceps (TM), lateral-triceps (TL), anterior deltoid (AD), posterior deltoid (PD), pectoralis major (PE), latissimus dorsi (LD)) were captured for a total of 640 trials per participant [25]. To enable the quantification of shared information across muscles with respect to specific task attributes, we formulated four discrete task parameters of length equal to the number of trials executed. These discrete variables represented the trial-to-trial variation in reaching direction (fwd vs bwd), speed (high vs low), and reaching target [P1-P8] (fig.3(A)).

In dataset 2 (fig.4(B)), 3 participants performed whole-body point-to-point reaching movements in various directions and to varying heights while EMG from 30 muscles (tibialis anterior, soleus, peroneus, gastrocnemius, vastus lateralis, rectus femoris, biceps femoris, gluteus maximus, erector spinae, pectoralis major, trapezius, anterior deltoid, posterior deltoid, biceps and triceps brachii) across both hemi-bodies were captured [26]. Like dataset 1, we formulated task parameters each representing a specific task attribute across trials (∼2160 trials per participant). In this case, we formed eight discrete task parameters representing start- and end-point target, -bar, and -height and both up-down (vertical) and left-right (horizontal) reaching directions.

Dataset 3 consisted of multiple trials from 17 participants performing level-ground walking, stair- and ramp-ascents/descents with various sub-conditions (walking speed, clockwise/counter-clockwise direction, different stair/ramp inclines etc.) (fig.4(C)) [27]. These locomotion modes were performed while 11 EMG signals were captured from the right lower-limb (Gluteus medius (GlutM), right external oblique (Obl), semitendinosus (ST), gracilis (GR), biceps femoris (BF), rectus femoris (RF), vastus lateralis (VL), vastus medialis (VM), soleus (SO), tibialis anterior (TA), gastrocnemius medialis (GM)), XYZ coordinates were captured bilaterally from 32 kinematic markers and 4 IMUs and a force-plate captured accelerations and dynamic features among the lower-limbs also. More detailed breakdowns of the experimental design for each dataset can be found at their parent publications [25–27].

For all datasets, we processed the EMG signals offline using a standardised approach [10]: the EMGs for each sample were digitally full-wave rectified, low-pass filtered (Butterworth filter; 20 Hz cut-off; zero-phase distortion), normalised to 1000 time-samples and then the signals were integrated over 20 time- step intervals yielding a waveform of ∼50 time-steps. To match the time-series lengths, we resampled the kinematic, dynamic and IMU recordings of dataset 3 using cubic-spline interpolation to match the EMG signals.

## References

[1] Macpherson T, Matsumoto M, Gomi H, Morimoto J, Uchibe E, Hikida T. Parallel and hierarchical neural mechanisms for adaptive and predictive behavioral control. Neural Networks. 2021 Dec 1;144:507–21.

[2] Kaplan HS, Thula OS, Khoss N, Zimmer M. Nested neuronal dynamics orchestrate a behavioral hierarchy across timescales. Neuron. 2020 Feb 5;105(3):562–76.

[3] Bruton M, O’Dwyer N. Synergies in coordination: a comprehensive overview of neural, computational, and behavioral approaches. Journal of Neurophysiology. 2018 Dec 1;120(6):2761–74.

[4] Bizzi E, Cheung VC. The neural origin of muscle synergies. Frontiers in computational neuroscience. 2013 Apr 29;7:51.

[5] Berret B, Delis I, Gaveau J, Jean F. Optimality and modularity in human movement: from optimal control to muscle synergies. InBiomechanics of anthropomorphic systems 2019 (pp. 105–133). Springer, Cham.

[6] Bernstein N. The co-ordination and regulation of movements. The co-ordination and regulation of movements. 1966.

[7] Latash ML. Synergy. Oxford University Press; 2008 Mar 18.

[8] Brenner N, Strong SP, Koberle R, Bialek W, Steveninck RR. Synergy in a neural code. Neural computation. 2000 Jul 1;12(7):1531–52.

[9] Turpin NA, Uriac S, Dalleau G. How to improve the muscle synergy analysis methodology?. European Journal of Applied Physiology. 2021 Apr;121(4):1009–25.

[10] d’Avella A, Lacquaniti F. Control of reaching movements by muscle synergy combinations. Frontiers in computational neuroscience. 2013 Apr 19;7:42.

[11] Ó’Reilly D, Delis I. A network information theoretic framework to characterise muscle synergies in space and time. Journal of Neural Engineering. 2022 Feb 18;19(1):016031.

[12] Alessandro C, Delis I, Nori F, Panzeri S, Berret B. Muscle synergies in neuroscience and robotics: from input-space to task-space perspectives. Frontiers in computational neuroscience. 2013 Apr 19;7:43.

[13] de Rugy A, Loeb GE, Carroll TJ. Are muscle synergies useful for neural control?. Frontiers in computational neuroscience. 2013 Mar 21;7:19.

[14] Krakauer JW, Ghazanfar AA, Gomez-Marin A, MacIver MA, Poeppel D. Neuroscience needs behavior: correcting a reductionist bias. Neuron. 2017 Feb 8;93(3):480–90.

[15] Cheung VC, Seki K. Approaches to revealing the neural basis of muscle synergies: a review and a critique. Journal of Neurophysiology. 2021 May 1;125(5):1580–97.

[16] Ronzano R, Lancelin C, Bhumbra GS, Brownstone RM, Beato M. Proximal and distal spinal neurons innervating multiple synergist and antagonist motor pools. Elife. 2021 Nov 2;10:e70858.

[17] Hug F, Del Vecchio A, Avrillon S, Farina D, Tucker K. Muscles from the same muscle group do not necessarily share common drive: evidence from the human triceps surae. Journal of Applied Physiology. 2021 Feb 1;130(2):342–54.

[18] Hug F, Avrillon S, Sarcher A, Del Vecchio A, Farina D. Correlation networks of spinal motor neurons that innervate lower limb muscles during a multi-joint isometric task. The Journal of Physiology. 2022 Jun 30.

[19] Alessandro C, Barroso FO, Prashara A, Tentler DP, Yeh HY, Tresch MC. Coordination amongst quadriceps muscles suggests neural regulation of internal joint stresses, not simplification of task performance. Proceedings of the National Academy of Sciences. 2020 Apr 7;117(14):8135–42.

[20] Valero-Cuevas FJ, Venkadesan M, Todorov E. Structured variability of muscle activations supports the minimal intervention principle of motor control. Journal of neurophysiology. 2009 Jul;102(1):59–68.

[21] Nazarpour K, Barnard A, Jackson A. Flexible cortical control of task-specific muscle synergies. Journal of Neuroscience. 2012 Sep 5;32(36):12349–60.

[22] Todorov E, Jordan M. A minimal intervention principle for coordinated movement. Advances in neural information processing systems. 2002;15.

[23] d’Avella A, Bizzi E. Shared and specific muscle synergies in natural motor behaviors. Proceedings of the national academy of sciences. 2005 Feb 22;102(8):3076–81.

[24] Tresch CV. MC Bizzi E 2005 Central and sensory contributions to the activation and organization of muscle synergies during natural motor behaviors. J Neurosci.;25:64196434.

[25] Delis I, Panzeri S, Pozzo T, Berret B. A unifying model of concurrent spatial and temporal modularity in muscle activity. Journal of neurophysiology. 2014 Feb 1;111(3):675–93.

[26] Hilt PM, Delis I, Pozzo T, Berret B. Space-by-time modular decomposition effectively describes whole-body muscle activity during upright reaching in various directions. Frontiers in computational neuroscience. 2018 Apr 3;12:20.

[27] Camargo J, Ramanathan A, Flanagan W, Young A. A comprehensive, open-source dataset of lower limb biomechanics in multiple conditions of stairs, ramps, and level-ground ambulation and transitions. Journal of Biomechanics. 2021 Apr 15;119:110320.

[28] Ince RA, Giordano BL, Kayser C, Rousselet GA, Gross J, Schyns PG. A statistical framework for neuroimaging data analysis based on mutual information estimated via a gaussian copula. Human brain mapping. 2017 Mar;38(3):1541–73.

[29] Blondel VD, Guillaume JL, Lambiotte R, Lefebvre E. Fast unfolding of communities in large networks. Journal of statistical mechanics: theory and experiment. 2008 Oct 9;2008(10):P10008.

[30] Mucha PJ, Richardson T, Macon K, Porter MA, Onnela JP. Community structure in time-dependent, multiscale, and multiplex networks. science. 2010 May 14;328(5980):876-8.

[31] Lancichinetti A, Fortunato S. Consensus clustering in complex networks. Scientific reports. 2012 Mar 27;2(1):1–7.

[32] Rubinov M, Sporns O. Complex network measures of brain connectivity: uses and interpretations. Neuroimage. 2010 Sep 1;52(3):1059–69.

[33] Averta G, Barontini F, Catrambone V, Haddadin S, Handjaras G, Held JP, Hu T, Jakubowitz E, Kanzler CM, Kühn J, Lambercy O. U-Limb: A multi-modal, multi-center database on arm motion control in healthy and post-stroke conditions. GigaScience. 2021 Jun;10(6):giab043.

[34] Funato T, Hattori N, Yozu A, An Q, Oya T, Shirafuji S, Jino A, Miura K, Martino G, Berger D, Miyai I. Muscle synergy analysis yields an efficient and physiologically relevant method of assessing stroke. Brain Communications. 2022 Aug 1;4(4):fcac200.

[35] Scano A, Mira RM, d’Avella A. Mixed matrix factorization: A novel algorithm for the extraction of kinematic-muscular synergies. Journal of Neurophysiology. 2022 Feb 1;127(2):529–47.

[36] Buongiorno D, Cascarano GD, Camardella C, De Feudis I, Frisoli A, Bevilacqua V. Task-oriented muscle synergy extraction using an autoencoder-based neural model. Information. 2020 Apr 17;11(4):219.

[37] Tresch MC, Saltiel P, Bizzi E. The construction of movement by the spinal cord. Nature neuroscience. 1999 Feb;2(2):162–7.

[38] Makarov SN, Noetscher GM, Nazarian A. Low-frequency electromagnetic modeling for electrical and biological systems using MATLAB. John Wiley & Sons; 2015 Jun 22.

[39] Benzi M, Klymko C. Total communicability as a centrality measure. Journal of Complex Networks. 2013 Dec 1;1(2):124–49.

[40] Clark DJ, Ting LH, Zajac FE, Neptune RR, Kautz SA. Merging of healthy motor modules predicts reduced locomotor performance and muscle coordination complexity post-stroke. Journal of neurophysiology. 2010 Feb;103(2):844–57.

[41] Schwartz MH, Rozumalski A, Steele KM. Dynamic motor control is associated with treatment outcomes for children with cerebral palsy. Developmental Medicine & Child Neurology. 2016 Nov;58(11):1139–45.

[42] Steele KM, Rozumalski A, Schwartz MH. Muscle synergies and complexity of neuromuscular control during gait in cerebral palsy. Developmental Medicine & Child Neurology. 2015 Dec;57(12):1176–82.

[43] Gallos LK, Makse HA, Sigman M. A small world of weak ties provides optimal global integration of self-similar modules in functional brain networks. Proceedings of the National Academy of Sciences. 2012 Feb 21;109(8):2825–30.

[44] Bunde A, Havlin S, editors. Fractals and disordered systems. Springer Science & Business Media; 2012 Dec 6.

[45] Bak P, Tang C, Wiesenfeld K. Self-organized criticality: An explanation of the 1/f noise. Physical review letters. 1987 Jul 27;59(4):381.

[46] Didier G, Valdeolivas A, Baudot A. Identifying communities from multiplex biological networks by randomized optimization of modularity. F1000Research. 2018;7.

[47] Newman ME, Girvan M. Finding and evaluating community structure in networks. Physical review E. 2004 Feb 26;69(2):026113.

[48] Magnani M, Hanteer O, Interdonato R, Rossi L, Tagarelli A. Community detection in multiplex networks. ACM Computing Surveys (CSUR). 2021 May 8;54(3):1–35.

[49] Kerkman JN, Daffertshofer A, Gollo LL, Breakspear M, Boonstra TW. Network structure of the human musculoskeletal system shapes neural interactions on multiple time scales. Science advances. 2018 Jun 27;4(6):eaat0497.

[50] Hardesty RL, Boots MT, Yakovenko S, Gritsenko V. Computational evidence for nonlinear feedforward modulation of fusimotor drive to antagonistic co-contracting muscles. Scientific reports. 2020 Jun 30;10(1):1–7.

[51] Cincotta M, Ziemann U. Neurophysiology of unimanual motor control and mirror movements. Clinical Neurophysiology. 2008 Apr 1;119(4):744–62.

[52] McGill W. Multivariate information transmission. Transactions of the IRE Professional Group on Information Theory. 1954 Sep;4(4):93–111.

[53] Schyns PG, Zhan J, Jack RE, Ince RA. Revealing the information contents of memory within the stimulus information representation framework. Philosophical Transactions of the Royal Society B. 2020 May 25;375(1799):20190705.

[54] Pickle NT, Grabowski AM, Auyang AG, Silverman AK. The functional roles of muscles during sloped walking. Journal of biomechanics. 2016 Oct 3;49(14):3244–51.

[55] Delis I, Hilt PM, Pozzo T, Panzeri S, Berret B. Deciphering the functional role of spatial and temporal muscle synergies in whole-body movements. Scientific reports. 2018 May 30;8(1):1–7.

[56] Brambilla C, Atzori M, Müller H, d’Avella A, Scano A. Spatial and temporal muscle synergies provide a dual characterization of low-dimensional and intermittent control of upper-limb movements. bioRxiv. 2022 Jan 1.

[57] Williams AH, Kim TH, Wang F, Vyas S, Ryu SI, Shenoy KV, Schnitzer M, Kolda TG, Ganguli S. Unsupervised discovery of demixed, low-dimensional neural dynamics across multiple timescales through tensor component analysis. Neuron. 2018 Jun 27;98(6):1099–115.

[58] Schneidman E, Bialek W, Berry MJ. Synergy, redundancy, and independence in population codes. Journal of Neuroscience. 2003 Dec 17;23(37):11539–53.

[59] Nigam S, Pojoga S, Dragoi V. Synergistic coding of visual information in columnar networks. Neuron. 2019 Oct 23;104(2):402–11.

[60] Luppi AI, Mediano PA, Rosas FE, Holland N, Fryer TD, O’Brien JT, Rowe JB, Menon DK, Bor D, Stamatakis EA. A synergistic core for human brain evolution and cognition. Nature Neuroscience. 2022 Jun;25(6):771–82.

[61] Urai AE, Doiron B, Leifer AM, Churchland AK. Large-scale neural recordings call for new insights to link brain and behavior. Nature neuroscience. 2022 Jan;25(1):11–9.

[62] Chiovetto E, Berret B, Delis I, Panzeri S, Pozzo T. Investigating reduction of dimensionality during single-joint elbow movements: a case study on muscle synergies. Frontiers in computational neuroscience. 2013 Feb 28;7:11.

[63] Del Vecchio A, Germer C, Kinfe TM, Nuccio S, Hug F, Eskofier B, Farina D, Enoka RM. Common synaptic inputs are not distributed homogeneously among the motor neurons that innervate synergistic muscles. BioRxiv. 2022 Jan 23:2022–01.

[64] Borzelli D, Vieira TD, Botter A, Gazzoni M, Lacquaniti F, d’Avella A. Independent synaptic inputs to motor neurons driving antagonist muscles. bioRxiv. 2022 Jan 1.

[65] Sartori L, Straulino E, Castiello U. How objects are grasped: the interplay between affordances and end-goals. PloS one. 2011 Sep 28;6(9):e25203.

[66] Dounskaia N, Shimansky Y, Ganter BK, Vidt ME. A simple joint control pattern dominates performance of unconstrained arm movements of daily living tasks. PloS one. 2020 Jul 13;15(7):e0235813.

[67] Santello M, Bianchi M, Gabiccini M, Ricciardi E, Salvietti G, Prattichizzo D, Ernst M, Moscatelli A, Jörntell H, Kappers AM, Kyriakopoulos K. Hand synergies: Integration of robotics and neuroscience for understanding the control of biological and artificial hands. Physics of life reviews. 2016 Jul 1;17:1–23.

[68] Dominijanni G, Shokur S, Salvietti G, Buehler S, Palmerini E, Rossi S, De Vignemont F, d’Avella A, Makin TR, Prattichizzo D, Micera S. The neural resource allocation problem when enhancing human bodies with extra robotic limbs. Nature Machine Intelligence. 2021 Oct;3(10):850–60.

[69] Delis I, Ince RA, Sajda P, Wang Q. Information-theoretic characterization of the neural mechanisms of active multisensory decision making. InInternational Conference on NeuroRehabilitation 2018 Oct 16 (pp. 584–588). Springer, Cham.

[70] Adams RA, Shipp S, Friston KJ. Predictions not commands: active inference in the motor system. Brain Structure and Function. 2013 May;218(3):611–43.

[71] Estrada E, Hatano N. Communicability in complex networks. Physical Review E. 2008 Mar 11;77(3):036111.

