## Supplementary materials for "Dissecting muscle synergies in the task space"

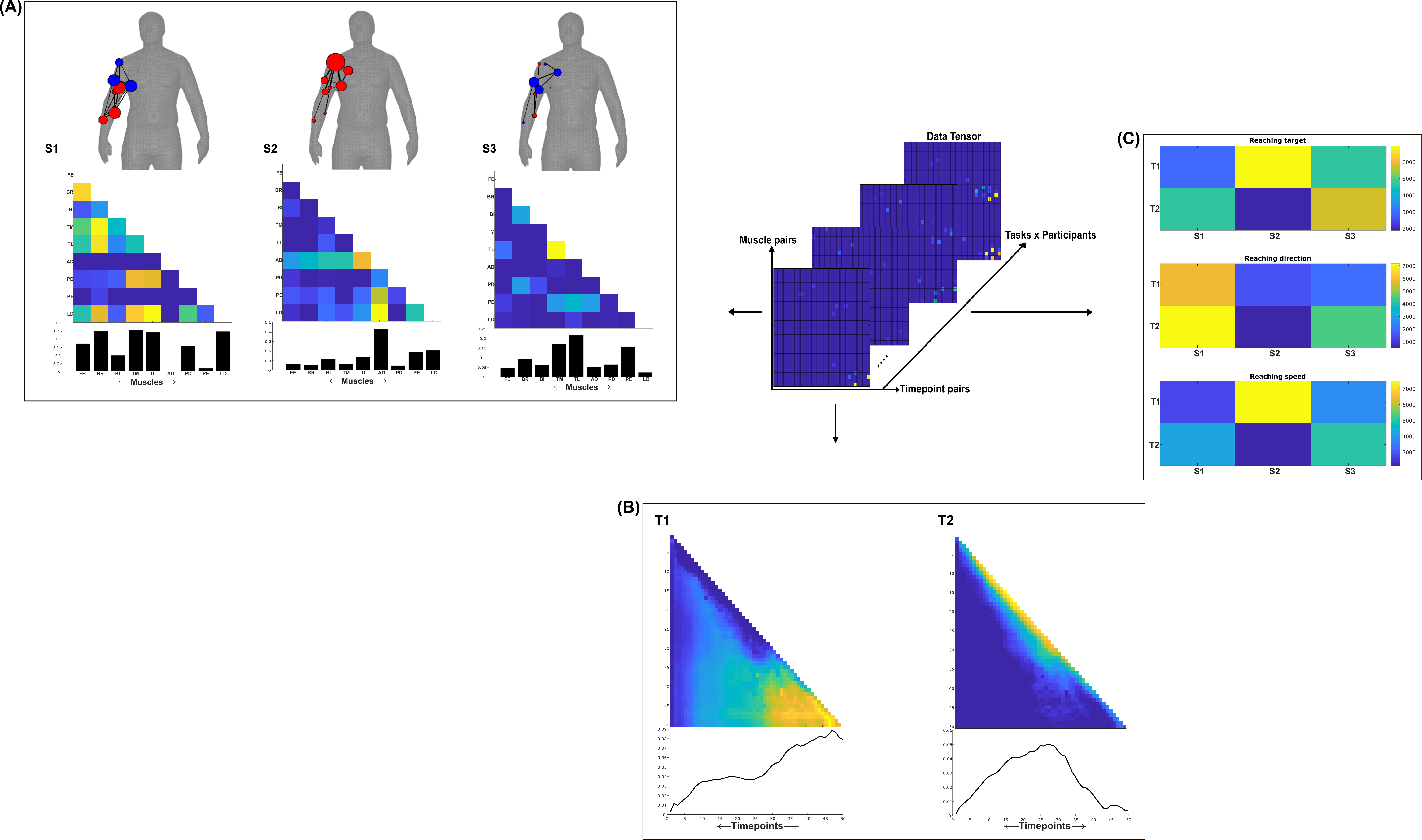

**Fig.1**: Three spatial (S1-S3) and two temporal task-irrelevant muscle networks (T1-T2) were empirically identified and extracted across participants and task parameters from dataset 1 using the NIF pipeline (Panel **A**-**B**) [10,23]. Human body models accompanying each spatial network illustrate their respective submodular structure with node colour and size and edge width indicating community affiliation [30], network centrality and connection strength respectively [35,43]. (Panel **C**) Activation coefficients are presented to the right of the networks, indicating their task parameter-specific scaling averaged across participants.

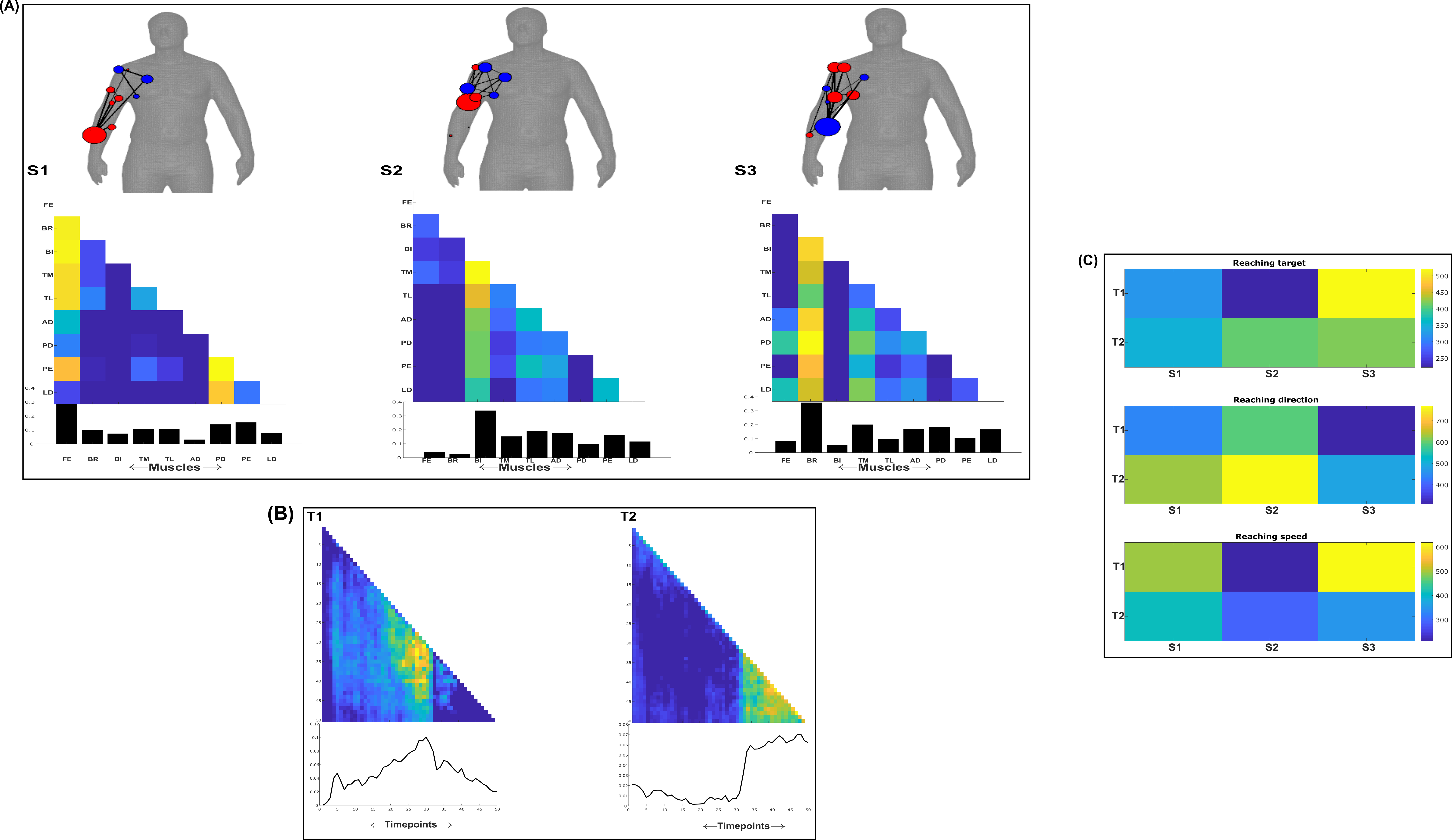

**Fig.2**: Three spatial (S1-S3) and two temporal task-redundant muscle networks (T1-T2) were empirically identified and extracted across participants and task parameters from dataset 1 using the NIF pipeline (Panel **A**-**B**) [10,23]. Human body models accompanying each spatial network illustrate their respective submodular structure with node colour and size and edge width indicating community affiliation [30], network centrality and connection strength respectively [35,43]. (Panel **C**) Activation coefficients are presented to the right of the networks, indicating their task parameter-specific scaling averaged across participants.

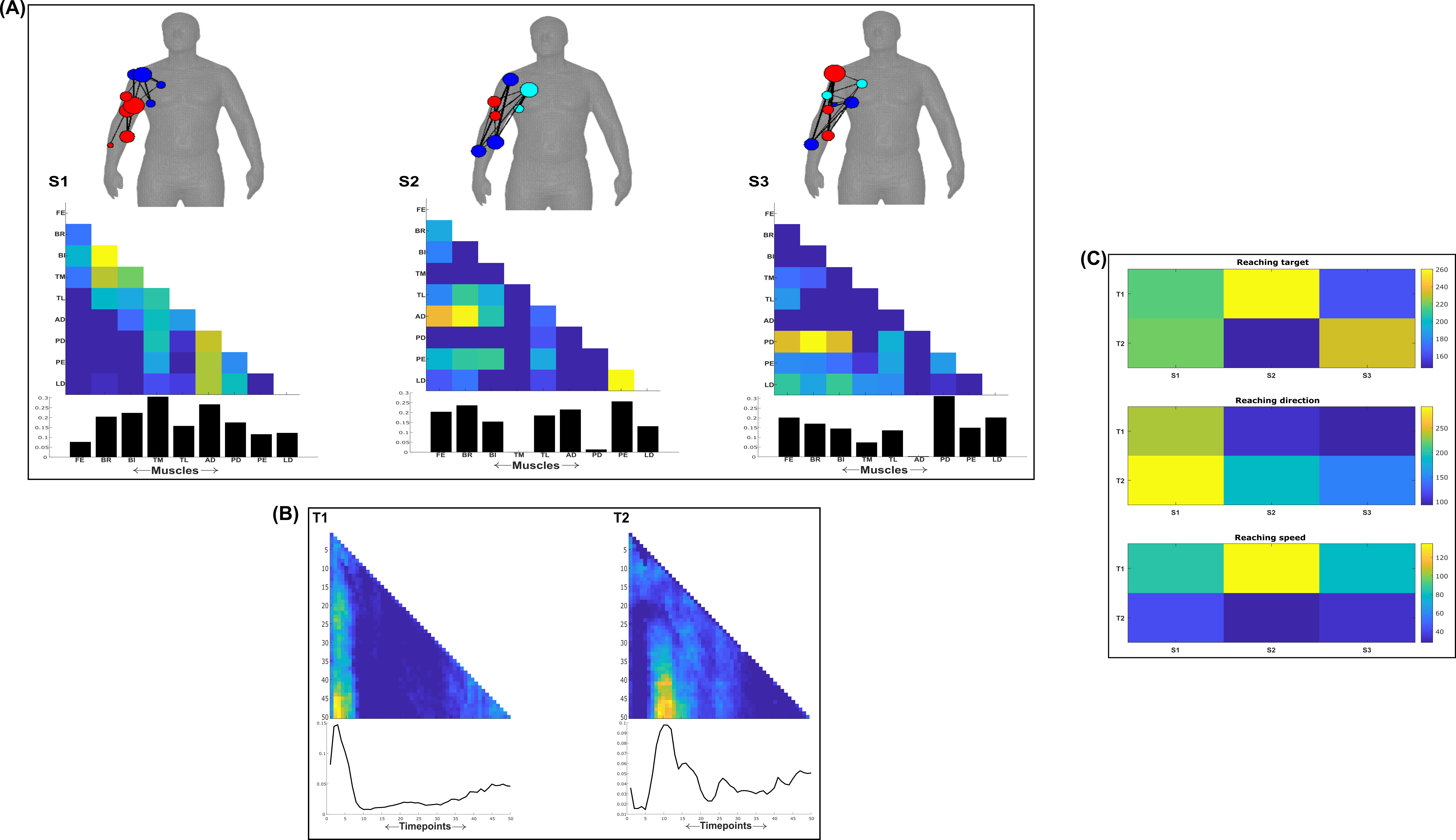

**Fig.3**: Three spatial (S1-S3) and two temporal task-synergistic muscle networks (T1-T2) were empirically identified and extracted across participants and task parameters from dataset 1 using the NIF pipeline (Panel **A**-**B**) [10,23]. Human body models accompanying each spatial network illustrate their respective submodular structure with node colour and size and edge width indicating community affiliation [30], network centrality and connection strength respectively [35,43]. (Panel **C**) Activation coefficients are presented to the right of the networks, indicating their task parameter-specific scaling averaged across participants.

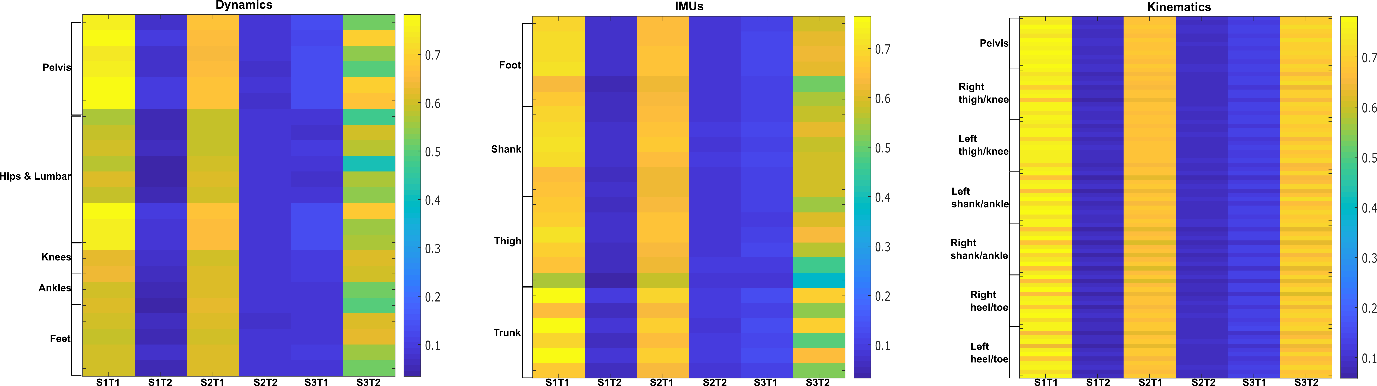

**Fig.4**: Task-irrelevant activation coefficients (Dataset 3) [25]. Dynamic, inertial motion unit (IMU) and kinematic data were captured from the bilateral lower-limbs while 17 participants performed various locomotion modes (i.e. stair ascents/descents, ramp inclines/declines and level-ground walking). Activation coefficients are averaged across participants.

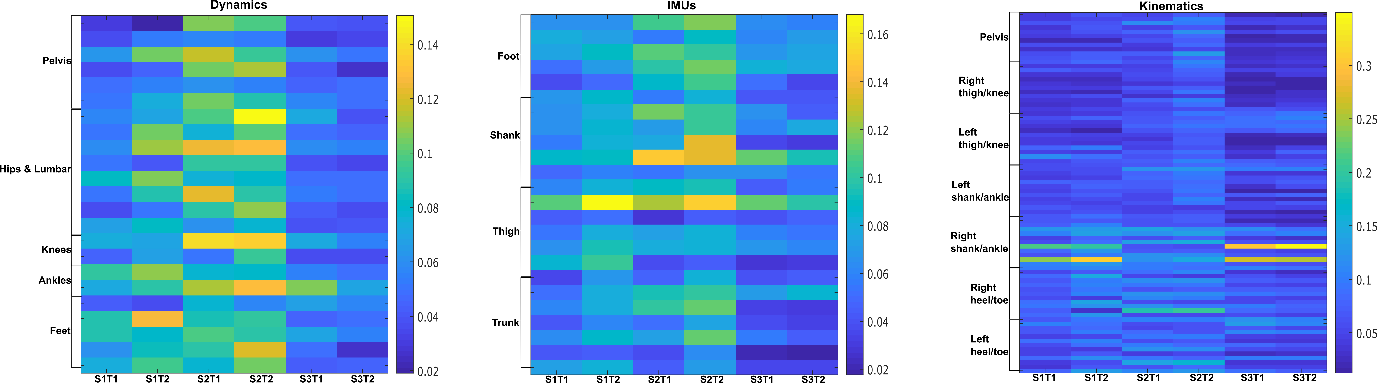

**Fig.5**: Task-redundant activation coefficients (Dataset 3) [25]. Dynamic, inertial motion unit (IMU) and kinematic data were captured from the bilateral lower-limbs while 17 participants performed various locomotion modes (i.e. stair ascents/descents, ramp inclines/declines and level-ground walking). Activation coefficients are averaged across participants.

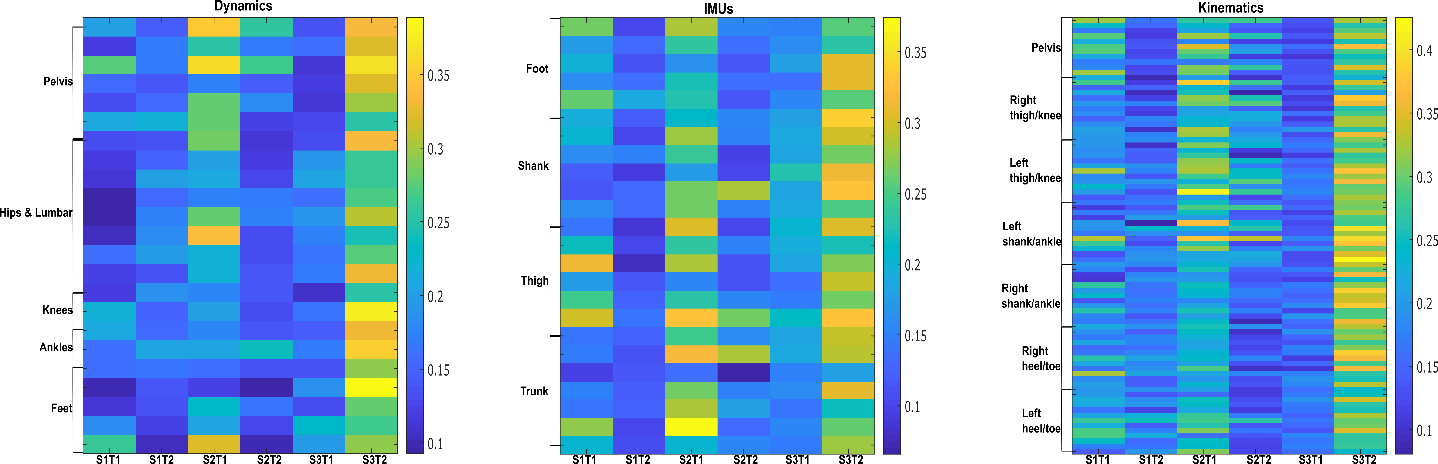

**Fig.6**: Task-synergistic activation coefficients (Dataset 3) [25]. Dynamic, inertial motion unit (IMU) and kinematic data were captured from the bilateral lower-limbs while 17 participants performed various locomotion modes (i.e. stair ascents/descents, ramp inclines/declines and level-ground walking). Activation coefficients are averaged across participants.
